# Tuft cell IL-17RB restrains IL-25 bioavailability and reveals context-dependent ILC2 hypoproliferation

**DOI:** 10.1101/2024.03.04.583299

**Authors:** Xiaogang Feng, Tilde Andersson, Julia Gschwend, Pascal Flüchter, Ivan Berest, Julian L. Muff, Daniele Carchidi, Antonie Lechner, Jeshua C. de Tenorio, Nina Brander, Ulrich Boehm, Christoph S. N. Klose, David Artis, Trese Leinders-Zufall, Frank Zufall, Christoph Schneider

## Abstract

The tuft cell–ILC2 circuit orchestrates rapid type 2 responses upon detecting microbe-derived succinate and luminal helminths. Our findings delineate key mechanistic steps, involving IP3R2 engagement and Ca^2+^ flux, governing IL-25 production by tuft cells triggered by succinate detection. While IL-17RB plays a pivotal intrinsic role in ILC2 activation, it exerts a regulatory function in tuft cells. Tuft cells exhibit constitutive *Il25* expression, placing them in an anticipatory state that facilitates rapid production of IL-25 protein for ILC2 activation. Tuft cell IL-17RB is crucial for restraining IL-25 bioavailability, preventing excessive tonic ILC2 stimulation due to basal *Il25* expression. Suboptimal ILC2 stimulation by IL-25 resulting from tuft cell *Il17rb*-deficiency or prolonged succinate exposure induces a state of hypoproliferation in ILC2s, also observed in chronic helminth infection. Our study offers critical insights into the regulatory dynamics of IL-25 in this circuit, highlighting the delicate tuning required for responses to diverse luminal states.

## Introduction

The single-layered epithelium of the small intestine separates luminal content from the underlying host tissue. In addition to their vital role in nutrient absorption, intestinal epithelial cells monitor the luminal status by detecting specific nutrients and microbes. These luminal signals can be sensed by distinct epithelial lineages and relayed to other cell types and tissues, leading to downstream responses that contribute to metabolic and immune regulation.

Chemosensory tuft cells are present in most mucosal epithelia, including the intestine, and have emerged as critical players in processing information regarding luminal status [1]. In the small intestine, tuft cells detect eukaryotic parasites such as helminths and protists, as well as certain shifts in bacterial composition and associated metabolites [2]. Upon helminth infection, tuft cells produce interleukin-25 (IL-25) and cysteinyl leukotrienes, activating lamina propria-resident group 2 innate lymphoid cells (ILC2) [3–6]. ILC2s respond by upregulating canonical type 2 cytokines, including IL-13, which promotes tuft and goblet cell differentiation. This tuft cell–ILC2 circuit is also activated by succinate which can be produced by *Tritrichomonas* protists or bacteria, stimulating tuft cells expressing the cognate G protein-coupled receptor (GPCR) SUCNR1 [7–9]. The acute succinate-mediated circuit activation depends on tuft cell-derived IL-25 for ILC2 activation, with cysteinyl leukotrienes being dispensable in that context [6–8]. Despite the constitutive expression of *Il25* transcript in tuft cells [3], the regulatory mechanisms governing IL-25 synthesis and release, for subsequent rapid ILC2 activation, remains unclear.

IL-25, previously known as IL-17E, is a member of the IL-17 cytokine family which contains the structurally related proteins IL-17A-F [10]. While best studied for IL-17A and F, this cytokine family is thought to engage canonical NF-κB signaling downstream of Act1 recruitment to a cytokine-specific multimeric receptor complex. The signaling-competent IL-25 receptor comprises two distinct chains, IL-17RB and IL-17RA [11]. Recent structural studies demonstrate that IL-25 binds specifically to IL-17RB units, allosterically initiating the formation of a ternary complex with the IL-17RA subunits, necessary for downstream signaling events [12]. ILC2s in the small intestine lamina propria express high levels of IL-17RB [7, 13]. Other immune and non-immune cells also reported to respond to IL-25 include dendritic cells and keratinocytes [14–16]. IL-25 is sufficient to induce type 2 cytokine expression in ILC2s, and this response is abolished in mice lacking *Il17rb* globally [17, 18]. However, selective genetic ablation of IL-25 signaling in ILC2s, to demonstrate its cell-intrinsic role under physiologically relevant stimulation of the tuft cell–ILC2 circuit, has not been reported so far. Notably, excessive homeostatic activation caused by IL-25 signaling in ILC2s is constrained by the ubiquitin-editing enzyme A20 (TNFAIP3) [7], as well as the cytokine-inducible SH2-containing protein (CISH) [19]. In contrast, little is known about the tuft cell-intrinsic regulation of circuit activity. Although tuft cells constitutively express the machinery for biosynthesis of IL-25 and cysteinyl leukotrienes, they do not strongly stimulate ILC2s in the absence of an agonistic signal [7, 8]. Thus, the mechanistic basis for how tuft cells control effector molecules, such as IL-25, remains to be defined.

In this study, we aimed to address important, yet unanswered questions pertaining to the regulation of IL-25 and the cell type-specific functions of its receptor, IL-17RB. We found that tuft cells rapidly release IL-25 triggered by intracellular calcium, activating ILC2s through the expression of IL-17RB on the latter. Tuft cell-intrinsic IL-17RB expression provides an unexpected mechanism for dampening homeostatic IL-25 release, protecting against the induction of a hypoproliferative state in ILC2s caused by prolonged exposure to IL-25.

## Results

### Succinate-induced IL-25 activates ILC2s in an ILC2-intrinsic, IL-17RB-dependent manner

To assess the ILC2-intrinsic requirement for IL-17RB, we generated mice with an *Il17rb* deficiency in *Il5*-expressing cells (*Il5*^R^;*Il17rb*^fl^) by crossing *Il17rb*^fl^ mice with “YRS” mice that express reporter alleles for ILC2 signature genes arginase-1 (Yarg; *Arg1*^YFP^), IL-5 (Red5; *Il5*^tdTomato-Cre^), and IL-13 (Smart13; *Il13*^hCD4^). In *Il5*^R/R^;*Il17rb*^fl/+^ control mice, we observed expression of IL-5 and IL-17RB in a majority of small intestinal ILC2s (Fig. 1a, b, S1a), consistent with prior literature [7]. CD4^+^ T cells from the same tissue showed neither substantial expression of IL-17RB, nor IL-5 (Fig. S1b). ILC2s from *Il5*^R/R^;*Il17rb*^fl/fl^ mice showed reduced IL-17RB staining; residual IL-17RB expression was found in IL-5^−/low^ ILC2s (Fig. 1a, b). IL-17RB expression was completely absent in ILC2s from *Il17rb*^−/−^ mice (Fig. 1a, b). We noted a slight reduction in Ki-67 expression in *Il17rb*-deficient ILC2s which was most pronounced in *Il17rb*^−/−^ mice, suggesting subtle tonic effects of IL-25 signaling in ILC2s (Fig. 1c). In these *Tritrichomonas*-free SOPF mice, some ILC2s expressed *Arg1*^YFP^ and they showed overall minimal expression of IL-13, in agreement with prior studies reporting low baseline ILC2 activation in the absence of luminal stimuli of the tuft cell–ILC2 circuit (Fig. S1c) [7]. To address the role of IL-17RB in ILC2s upon circuit activation, we treated mice with the luminal tuft cell agonist succinate in drinking water. After four days, control mice readily upregulated expression of IL-13 and Ki-67, which was significantly reduced in *Il5*^R/+^;*Il17rb*^fl/fl^ mice and fully abrogated in *Il17rb*^−/−^ mice. (Fig. 1d–e, S1d). To further establish the ILC2-intrinsic requirement for IL-17RB, we crossed *Il17rb*^fl/fl^ to *Nmur1*^iCre^ mice [20, 21]. *Nmur1*^iCre^*;Il17rb*^fl/fl^ mice showed near complete *Il17rb* deletion in ILC2s compared to littermate controls (Fig. 1f). Furthermore, expression of IL-13 and Ki-67 in ILC2 was abrogated in *Nmur1*^iCre^*;Il17rb*^fl/fl^ mice that were treated with succinate (Fig. 1h-i). These results firmly established that IL-25 directly stimulates ILC2s through their IL-17RB following activation of tuft cells with succinate.

**Figure 1.**
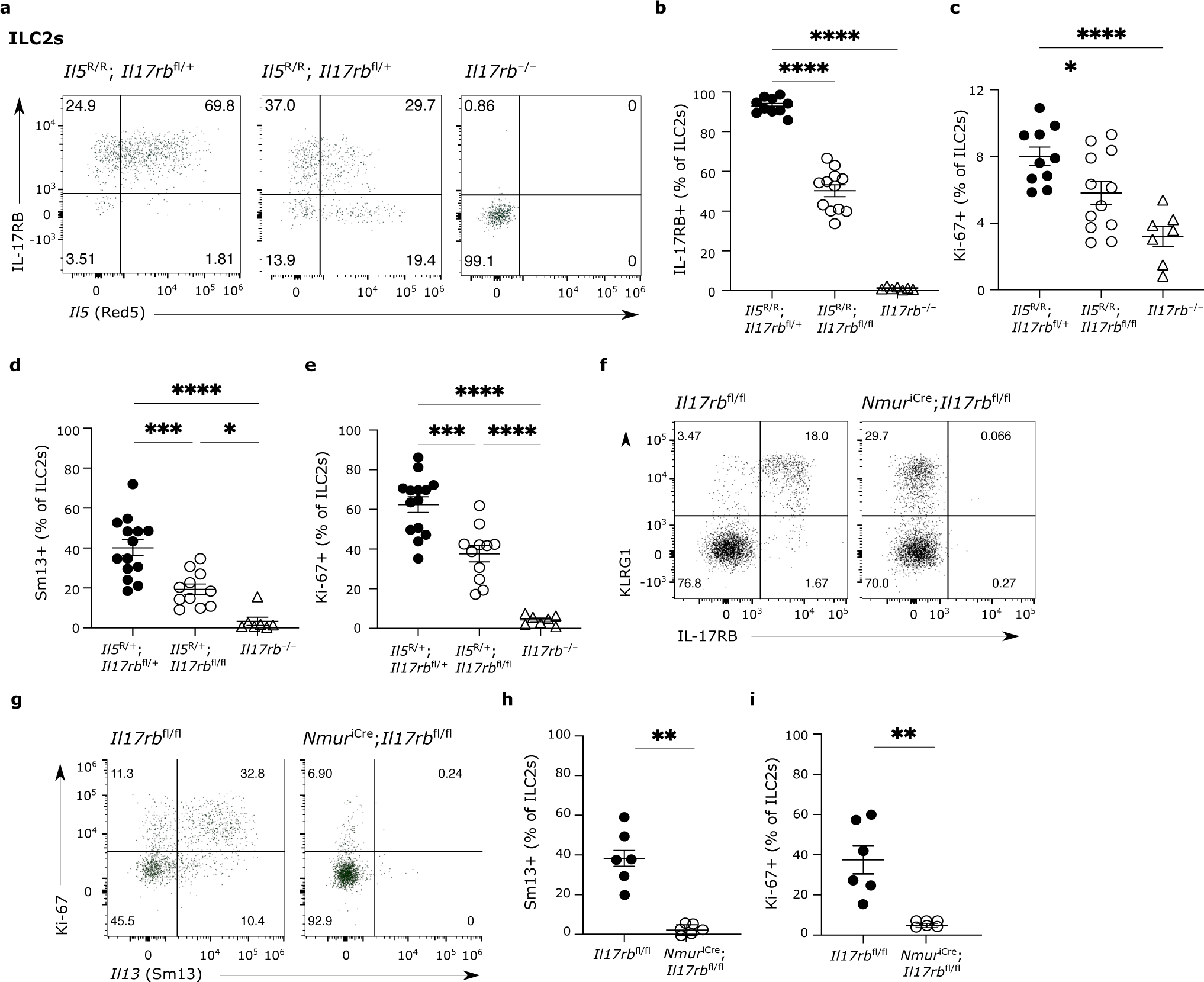
Succinate-induced IL-25 activates ILC2s in an ILC2-intrinsic, IL-17RB-dependent manner. (a-c) The small intestine (SI) of naïve *Il5*^R/R^;*Il17rb*^fl/+^, *Il5*^R/R^;*Il17rb*^fl/fl^, and *Il17rb*^−/−^ (*Il5*^+/+^) mice was analyzed. (a) Expression of IL-17RB and *Il5*-tdTomato (Red5) reporter by ILC2s. (b,c) Frequencies of IL-17RB^+^ (b) and Ki-67^+^ (c) ILC2s were quantified by flow cytometry. (d,e) *Il5*^R/+^;*Il17rb*^fl/+^, *Il5*^R/+^;*Il17rb*^fl/fl^, and *Il17rb*^−/−^ mice were treated with succinate for 4 days and the percentage of IL-13 (Sm13)^+^ (d) and Ki-67^+^ (e) ILC2s in the SI was quantified by flow cytometry. (f-i) *Nmur1*^iCre^;*Il17rb*^fl/fl^ and *Il17rb*^fl/fl^ littermate mice were treated with succinate for 4 days and the SI analyzed. (f) Expression of KLRG1 and IL-17RB by Lin^−^ cells. (g) Expression of Ki-67^+^ and IL-13 (Sm13)^+^ reporter by ILC2s. Frequencies of IL-13 (Sm13)^+^ (h) and Ki-67^+^ (i) ILC2s were quantified by flow cytometry. (a-c, f-i) Data is pooled from 2 independent experiments. (d,e) Data is pooled from 3 independent experiments. *, P 0.01–0.05; **, P 0.01–0.001; ****, P < 0.0001.

### Succinate-induced tuft cell activation depends on IP3R2-mediated cytosolic Ca^2+^ activity

Intrigued by the capacity of the succinate–tuft cell axis to rapidly stimulate ILC2s via IL-25 signaling, we further explored the mechanism underlying this response. Comparing expression of an *Il25*-tdTomato transcriptional reporter allele between tuft cells of mice treated with water or succinate revealed no marked alteration in *Il25* transcript, nor in the frequency of *Il25*-tdTomato^+^ cells (Fig. 2a-c, S2a). These results are in agreement with prior findings which demonstrated constitutive expression of *Il25* mRNA in tuft cells of various tissues [3]. Tuft cells engage canonical taste transduction signaling components when triggered by luminal agonists such as succinate, including the Ca^2+^-activated monovalent cation channel TRPM5 [7–9]. We therefore hypothesized that increasing cytosolic Ca^2+^ is a critical step in tuft cells for their production of IL-25. To test this, we visualized Ca^2+^ activity in isolated villi using the genetically encoded fast Ca^2+^ sensor GCaMP6f, expressed in TRPM5^+^ tuft cells, as recently published for tracheal tuft cells [22, 23]. In these *Trpm5*^Cre^;*R26*^GCaMP6f^ mice, temporally and spatially resolved Ca^2+^ signals can be recorded by confocal imaging in tuft cells in their native cellular environment in small intestinal villus epithelium (Fig. 2d, S2b,c). Exposure to succinate produced reproducible and concentration-dependent transient Ca^2+^ elevations in TRPM5^+^ tuft cells (Fig. 2e-g, Suppl. movie 1). The succinate-induced Ca^2+^ response was abolished when sarcoplasmic/endoplasmic reticulum calcium adenosine triphosphatase (ATPase) (SERCA) was inhibited with cyclopiazonic acid (CPA), indicating that Ca^2+^ release from intracellular Ca^2+^ stores is required for these responses (Fig. 2h,i). Type II taste bud cells release the second messenger ATP in a process that depends on the calcium channel inositol 1,4,5-trisphosphate receptor 3 (IP3R3) [24]. However, when analyzing RNA-sequencing data of small intestinal epithelial cells, we noted that tuft cells differentially express high levels of *Itpr2* transcript (encoding IP3R2) compared to non-tuft epithelial cells, whereas *Itpr3* and *Itpr1* transcripts were expressed at lower levels and comparable to those in other epithelial cells (Fig. 2j). Using a Cal630 Ca^2+^ indicator dye, we then measured Ca^2+^ responses in tuft cells from *Il25*^tdTomato^;*Itpr2*^+/−^ and *Il25*^tdTomato^;*Itpr2*^−/−^ mice (Fig. 2k, S2d). This analysis revealed a strongly impaired response to succinate in IP3R2-deficient tuft cells (Fig. 2l-n). Overall, our data suggested that IL-25 is controlled post-transcriptionally via IP3R2-regulated cytosolic Ca^2+^ activity.

**Figure 2.**
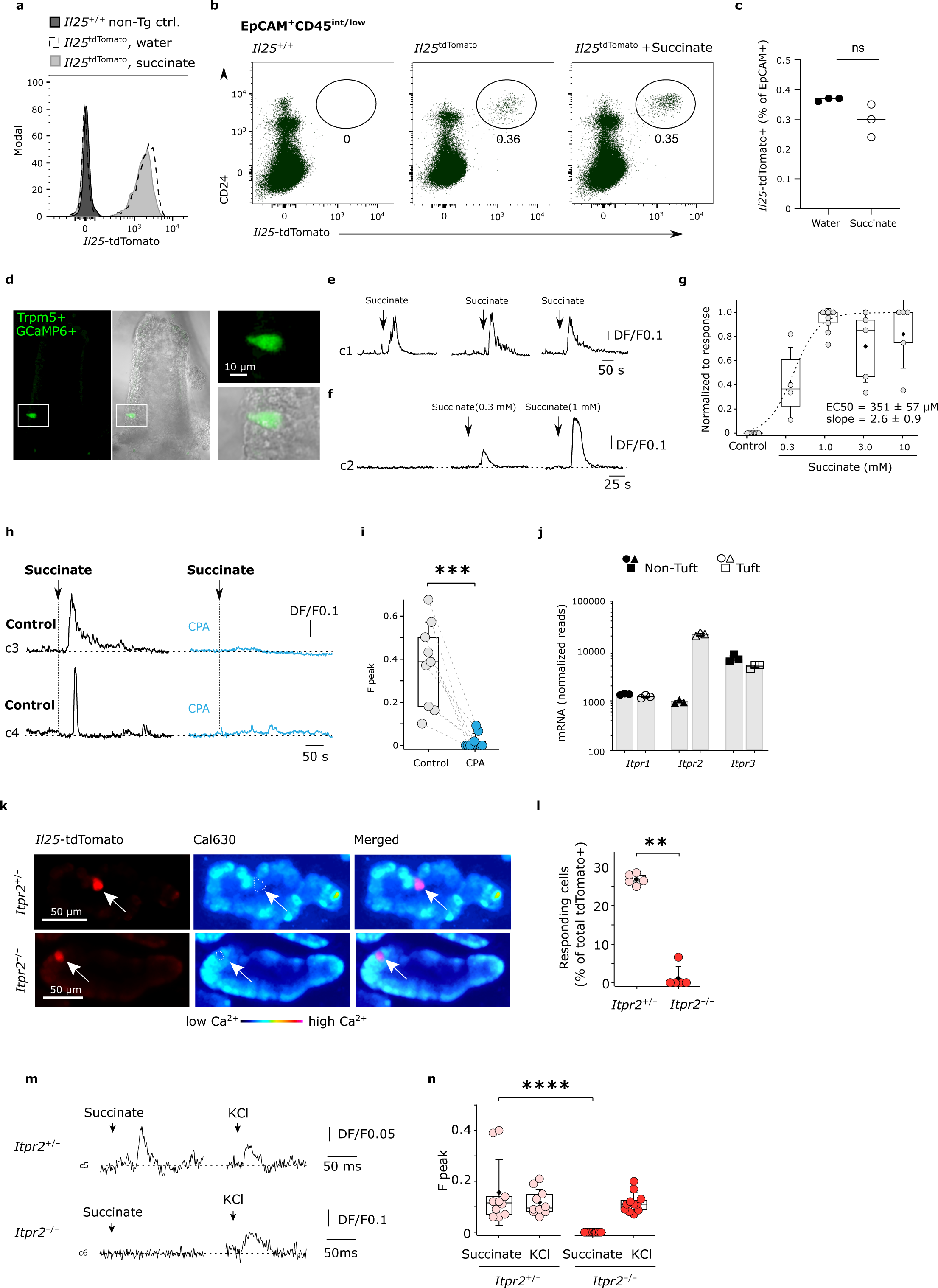
Succinate-induced tuft cell activation depends on IP3R2-mediated cytosolic Ca2^+^ activity. (a-c) Mice encoding the *Il25*^tdTomato^ reporter (Flare25) were treated with succinate or regular drinking water, and the small intestine (SI) was analyzed on day 4. (a) Expression of *Il25*^tdTomato^ in CD24^+^EpCAM^+^ tuft cells was determined by flow cytometry analysis and compared to tuft cells from a reporter-negative mouse. Expression and frequencies of *Il25*^tdTomato+^CD24^+^ tuft cells gated on EpCAM^+^CD45^int/low^ cells are shown (b) and the percentage of *Il25*^tdTomato+^cells was quantified (c). (d-i) Tuft cell Ca^2+^ activity was imaged using *Trpm5*^Cre^;*R26*^GCaMP6f^ mice. (d) Tuft cells (*green*) can be identified in an *ex vivo* scraped villi preparation from the ileum. A GCaMP6f-expressing tuft cell at the lower end of a villus (white box) has been magnified. (e) Reproducible Ca^2+^ response time courses of a tuft cell induced by 1 mM succinate. (f) Ca^2+^ response time courses of a tuft cell to increasing concentrations of succinate. (g) Dose response curve of Ca^2+^ peak responses normalized to the maximum response of an individual tuft cell. (h) Example traces of two tuft cells stimulated with 1 mM succinate before (control) and after 10 μM CPA treatment (blue). (i) Box plot summarizing the peak Ca^2+^ responses from preparations as in (h). (Fpeak) in 9 tuft cells, 3 mice. Paired t-test. (j) mRNA expression of *Itpr1*, *Itpr2*, and *Itpr3* in tuft cells and non-tuft epithelial cells from the SI, analyzed from dataset published by Nadjsombati *et al*.. (k-n) Tuft cell Ca^2+^ activity with a Cal630 indicator dye in ileum villi preparations from *Il25*^tdTomato^ (Flare25) *Itpr2*^+/−^ and *Itpr2*^−/−^ mice. (k-n) Confocal image visualizing tuft cells based on their *Il25*^tdTomato^ (Flare25) expression (*red*) after loading with the Cal630 indicator. (k) The merged image indicates that the tuft cells contain the Ca^2+^ indicator dye. (l) The percentage of succinate-responding tuft cells was quantified (5 mice/genotype; Mann-Whitney test). (m and n) Examples (M) of Ca^2+^ response time courses and quantification of peak response (n) to 1 mM succinate and 60 mM KCl (positive control) in tuft cells from an *Il25*^tdTomato^;*Itpr2*^+/−^ and *Il25*^tdTomato^;*Itpr2*^−/−^ mouse. Data are from one experiment representative of at least two independent experiments (a-f, h, k, m) or pooled from independent experiments (g, i, l, n). P values indicated; ns, P ≥ 0.05. Box plot in (n) summarizing the Fpeak values of independent tuft cell measurements: 5 mice/genotype; Mann-Whitney test; data are expressed as mean ± SD; box plots display the interquartile (25 - 75%) ranges, median (line) and mean (black square) values with whiskers indicating SD values.

### Succinate-elicited IL-25 production in tuft cells is triggered by IP3R2

To assess *in vivo* whether post-transcriptionally regulated IL-25 production, downstream of succinate receptor stimulation, is controlled by IP3R2-mediated Ca^2+^ activity, we stimulated *Itpr2*^−/−^ mice with succinate drinking water. Indeed, succinate-induced expression of IL-13 and Ki-67 by ILC2 was abrogated in *Itpr2*^−/−^ compared to *Itpr2*^+/−^ littermate controls (Fig. 3a,b). Notably, tuft cells from both genotypes expressed comparable levels of *Il25* transcript as indicated by the *Il25*-tdTomato reporter signal (Fig. 3c). When activation of tuft cells was bypassed by directly injecting rIL-25, ILC2s of *Itpr2*^−/−^ mice responded comparably well to those of *Itpr2*^+/−^ littermates, demonstrating that IP3R2 is not required in ILC2 for their stimulation by IL-25 (Fig. 3d,e). To address whether IL-25 protein is indeed released, binding to IL-17RB and activating ILC2s following tuft cell stimulation with succinate, we injected wild-type mice with an anti-IL-17RB blocking antibody [25] during the days of succinate treatment (Fig. 3f). Such acute blockade of IL-17RB potently prevented activation of ILC2s (Fig. 3g,h). Altogether, these results show that an IP3R2-dependent cytosolic Ca^2+^ signaling axis mediates post-transcription control of IL-25, triggering production and release when tuft cells are stimulated with succinate.

**Figure 3.**
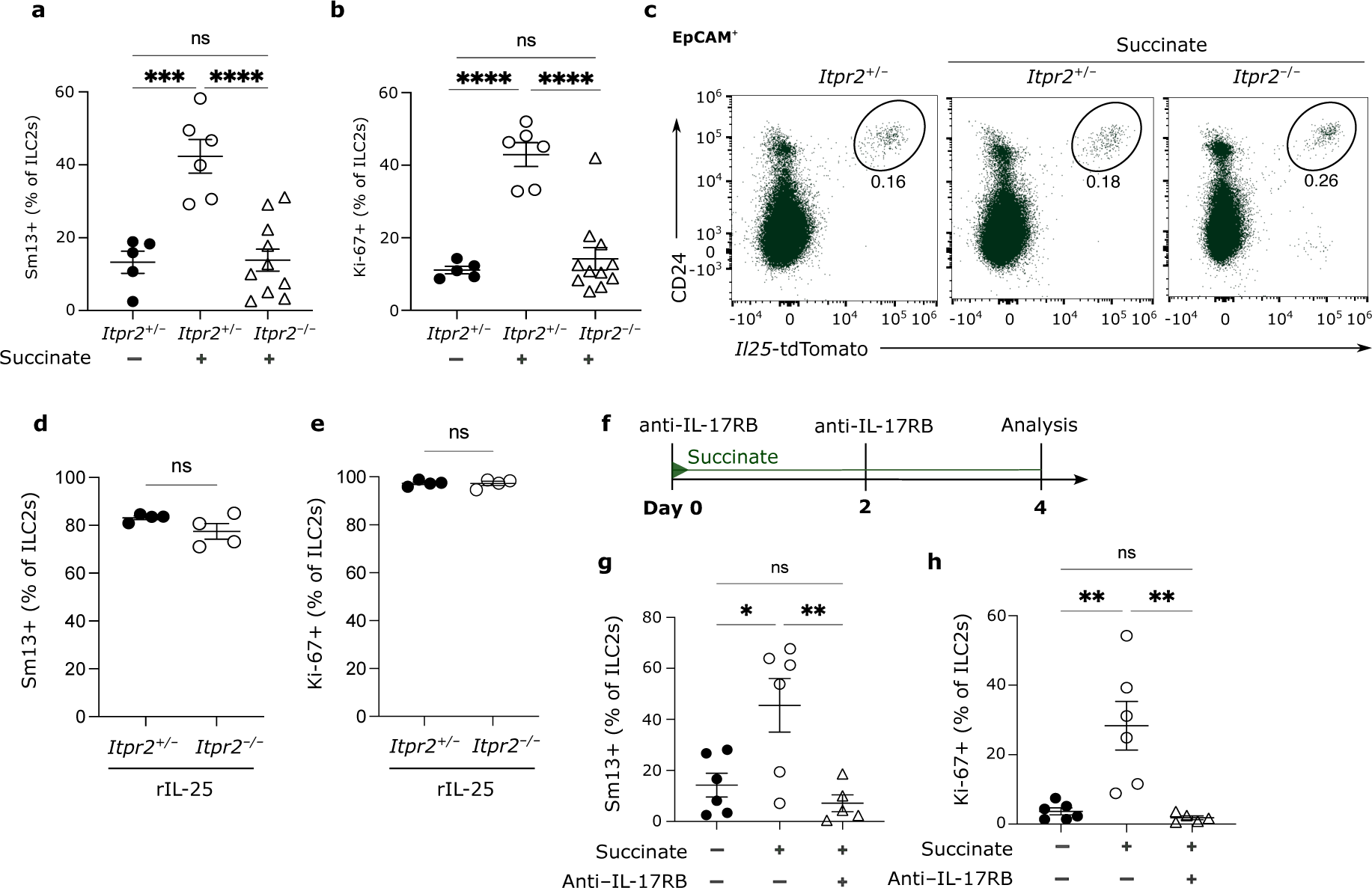
Succinate-elicited IL-25 production in tuft cells is triggered by IP3R2. (a-c) *Il25*^tdTomato^;*Itpr2*^−/−^ and *Il25*^tdTomato^;*Itpr2*^+/−^ littermate mice were treated with succinate and the small intestine was analyzed 4 days later. Frequencies of IL-13 (Sm13)^+^ (a) and Ki-67^+^ (b) ILC2s were quantified by flow cytometry. (c) Flow cytometry analysis of *Il25*^tdTomato^ (Flare25) reporter and CD24^+^ expression, gated on EpCAM^+^ cells. (d,e) *Il25*^tdTomato^;*Itpr2*^−/−^ and *Il25*^tdTomato^;*Itpr2*^+/−^ littermate mice were treated with 1 μg rIL-25Fc on 3 consecutive days and the small intestine was analyzed on the day after the last injection. Frequencies of IL-13 (Sm13)^+^ (d) and Ki-67^+^ (e) ILC2s were quantified by flow cytometry. (f-h) Mice expressing the IL-13 (Sm13) reporter were treated with succinate for 4 days and injected with PBS or anti-IL-17RB blocking antibody on d0 and d2, according to the schematic in (f). Frequencies of IL-13 (Sm13)^+^ (g) and Ki-67^+^ (h) ILC2s were quantified by flow cytometry. (a-e) Data pooled from 2 independent experiments. (d,e,g,h) Data from one experiment, representative of 3 independent experiments.*, P 0.01–0.05; **, P 0.01–0.001; ****, P < 0.0001.

### Tuft cell IL-17RB controls microbiota-independent ILC2 activation and restrains homeostatic circuit activation

We and others had noted the distinct expression of *Il17rb* mRNA in tuft cells compared to other intestinal epithelial cells of mice and humans [8, 26]. We also detected IL-17RB expression in tuft cells on the protein level (Fig. 4a). This unusual co-expression of the ligand-receptor pair, IL-25 and IL-17RB, prompted us to test the role of IL-17RB in regulation of tuft cell IL-25. For this we generated *Vil1*^Cre^*;Il17rb*^fl/fl^ mice which delete *Il17rb* in intestinal epithelial cells. Absence of IL-17RB staining was confirmed in tuft cells of *Vil1*^Cre^;*Il17rb*^fl/fl^ and *Il17rb*^−/−^ mice (Fig. 4a). To our surprise, tuft cell numbers were increased in the small intestine of *Vil1*^Cre^;*Il17rb*^fl/fl^ (Fig 4b,c). The concomitant increase in goblet cells indicated an elevated basal activity of the tuft cell–ILC2 circuit in *Vil1*^Cre^;*Il17rb*^fl/fl^ mice (Fig. 4d). Indeed, IL-13 expression by small intestinal ILC2s of *Vil1*^Cre^;*Il17rb*^fl/fl^ mice was significantly increased compared to ILC2s from their *Il17rb*^fl/fl^ littermates, while their Ki-67 expression did not noticeably differ (Fig. 4e-g). IL-13 expression was predominantly found in a compartment of *Arg1*-YFP^−^IL-17RB^+^ ILC2s, a phenotype that has been previously described for IL-25-stimulated ILC2s of the small intestinal lamina propria [7, 27] (Fig. S3a,b). A comparison of ILC2s from *Vil1*^Cre^;*Il17rb*^fl/fl^ and *Il17rb*^fl/fl^ littermates by RNA-sequencing showed similar expression in ILC2 signature genes, while expression of markers associated with ILC2 activation were increased in ILC2s from *Vil1*^Cre^;*Il17rb*^fl/fl^ mice (Fig. 4h,i).

**Figure 4.**
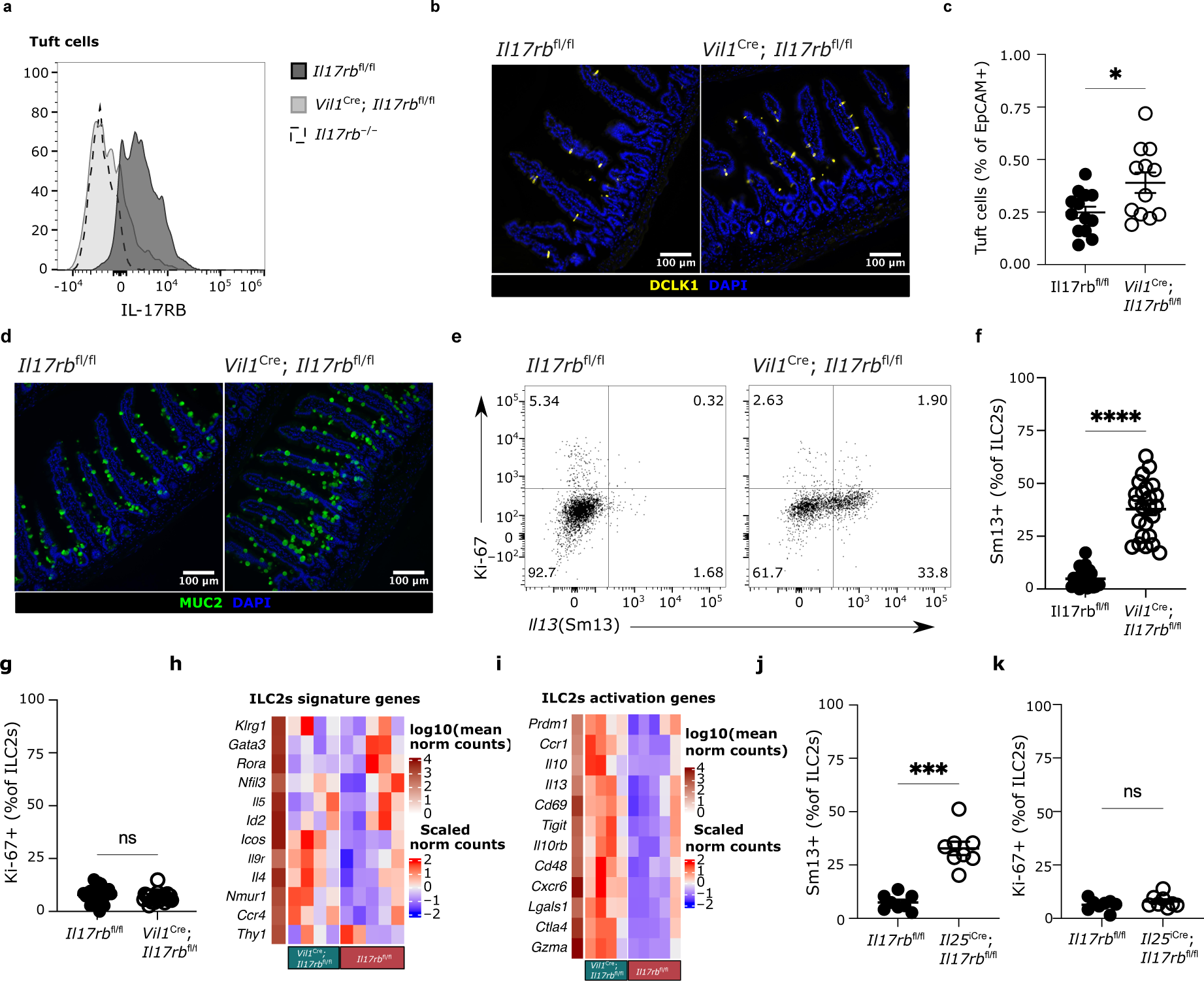
Tuft cell IL-17RB restrains homeostatic circuit activation. (a-g) The small intestine (SI) of naïve *Vil1*^Cre^;*Il17rb*^fl/fl^ and *Il17rb*^fl/fl^ littermate mice was analyzed. (a) Expression of IL-17RB in tuft cells was measured by flow cytometry and compared to that in *Il17rb*^−/−^ mice. (b) Representative immunofluorescence picture of DCLK1 (yellow) in SI (scale bars, 100 μm). (c) Quantification of tuft cells by flow cytometry. (d) Representative immunofluorescence picture of MUC2 (green) in SI (scale bars, 100 μm). (e) Flow cytometry analysis of Ki-67 and IL-13 (Sm13)-reporter expression by ILC2s. (f, g) Frequencies of IL-13 (Sm13)^+^ (f) and Ki-67^+^ (g) ILC2s, quantified by flow cytometry. (h, i) Heatmaps showing relative expression of ILC2 signature genes (h) and genes associated with ILC2 activation (i), generated from bulk RNA sequencing of FACSorted ILC2s (CD45^+^Lin^−^KLRG1^+^) from SI of *Vil1*^Cre^;*Il17rb*^fl/fl^ and *Il17rb*^fl/fl^ mice. (j, k) The SI of naïve *Il25*^iCre/+^;*Il17rb*^fl/fl^ and *Il17rb*^fl/fl^ littermate mice was analyzed. Frequencies of IL-13 (Sm13)^+^ (j) and Ki-67^+^ (k) ILC2s, quantified by flow cytometry. (a-g) Data representative of, and pooled from, 2 independent experiments. (h,i) Data from one experiment. (j,k) Data pooled from 2 independent experiments.*, P 0.01–0.05; **, P 0.01–0.001; ****, P < 0.0001.

To confirm that the elevated tonic activation of ILC2s in *Vil1*^Cre^;*Il17rb*^fl/fl^ mice was a result of *Il17rb*-deficiency in tuft cells, we also generated *Il25*^iCre/+^;*Il17rb*^fl/fl^ animals, in which deletion is limited to tuft cells due to their unique expression of *Il25* [3, 28]. Though recombination efficiency proved lower as compared to the *Vil1*^Cre^;*Il17rb*^fl/fl^ mice, IL-17RB expression was depleted in a majority of tuft cells of *Il25*^iCre/+^;*Il17rb*^fl/fl^ mice, resulting in a subtle increase in tuft cell abundance (Fig. S3c,d). In addition, ILC2s showed higher IL-13 expression, with no observed change in Ki-67, consistent with the elevated activation of ILC2s observed in *Vil1*^Cre^;*Il17rb*^fl/fl^ mice (Fig. 4j,k).

IL-25^+^ tuft cells emerge in the epithelium of the mouse small intestine shortly before weaning [7]. To test whether tuft cell IL-17RB controls ILC2 activity already at such an early age, we assessed the expression of IL-13 in ILC2s from 1-and 3-week-old mice. Consistent with the low abundance of tuft cells, IL-13 expression was low at the age of 1 week and did not differ between *Il17rb*^fl/fl^ and *Vil1*^Cre^;*Il17rb*^fl/fl^ mice (Fig. 5a,b). At 3 weeks of age, ILC2s from *Vil1*^Cre^;*Il17rb*^fl/fl^ mice showed an elevated activation state marked by a significant increase in the expression of IL-13 and KLRG1 (Fig. 5c-e), which was further confirmed in *Il25*^iCre/+^;*Il17rb*^fl/fl^ mice (Fig. 5f-i). Since our animals were free of *Tritrichomonas* protists, a major driver of ‘homeostatic’ circuit activation [4, 7], we wanted to determine if ILC2 activation in mice with tuft cell IL-17RB deficiency is driven by other intestinal microbes. However, *Vil1*^Cre^;*Il17rb*^fl/fl^ offspring still showed the same increase in IL-13 and KLRG1 expression compared to their littermate controls when treated before and after birth with a cocktail of antibiotics to deplete the microbiota (Fig. 5j-m). Collectively, these results suggested that tuft cell-intrinsic IL-17RB regulates microbiota-independent ILC2 activation, a process that commences during a postnatal stage when both cell types emerge in the small intestine, and then persist into adulthood.

**Figure 5.**
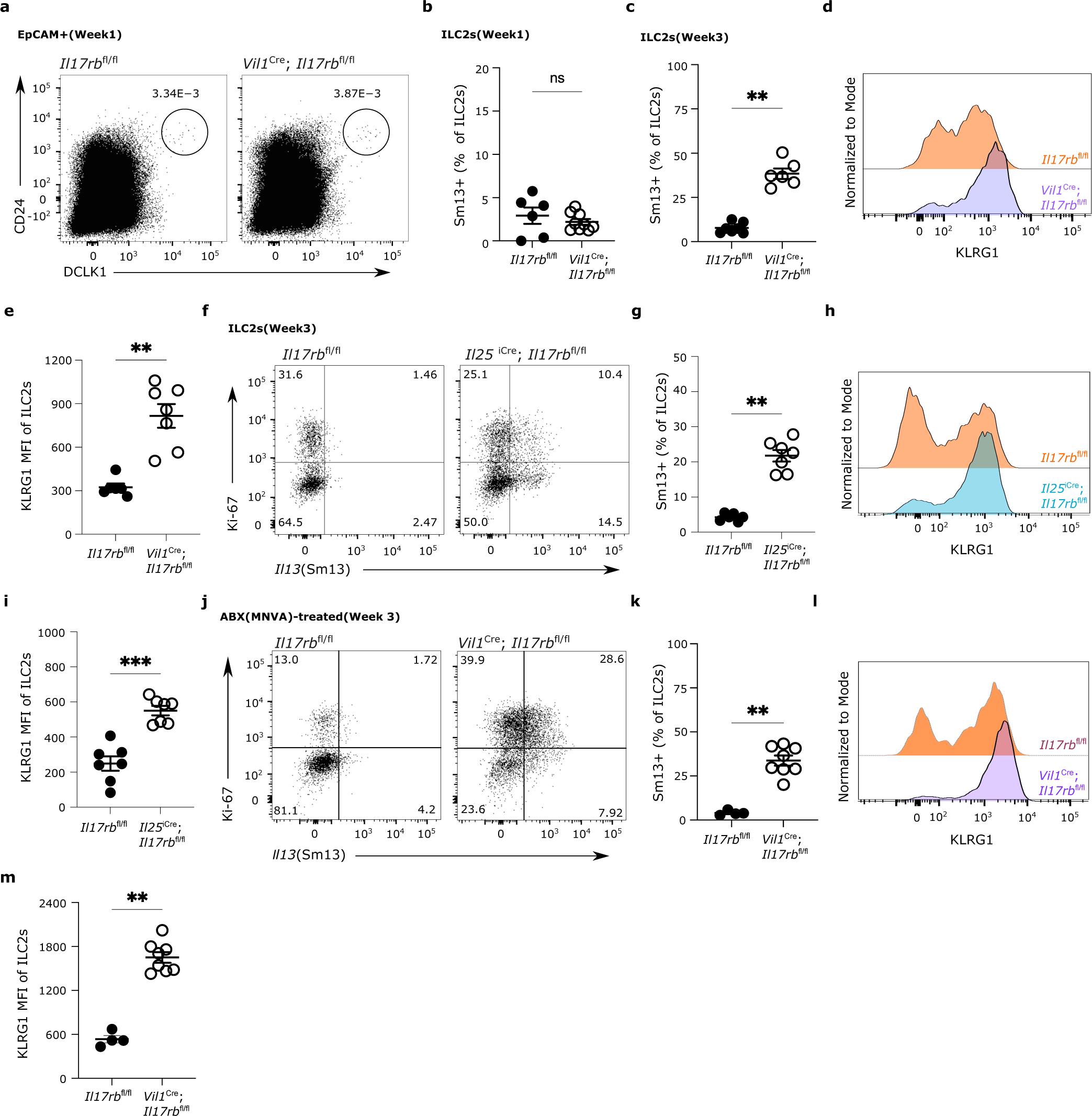
Tuft cell-intrinsic IL-17RB controls microbiota-independent ILC2 activation in young mice. (a-e) The small intestine (SI) of *Vil1*^Cre^;*Il17rb*^fl/fl^ and *Il17rb*^fl/fl^ littermate mice was analyzed at the age of one (a,b) or three (c-e) weeks. (a) Flow cytometry analysis of tuft cells, gated as DCLK1^+^CD24^+^ cells of EpCAM^+^. (b,c) Frequencies of IL-13 (Sm13)^+^ ILC2s at one (b) or 3 (c) weeks of age, quantified by flow cytometry. (d,e) Expression (d) and quantification (e) of KLRG1 in ILC2s, analyzed by flow cytometry. (f-i) Analysis of SI from 3-week-old *Il25*^iCre/+^;*Il17rb*^fl/fl^ and *Il17rb*^fl/fl^ littermate mice. (f,g) Flow cytometry analysis of Ki-67 and Sm13 reporter expression by ILC2s (f) and quantification of IL-13 (Sm13)^+^ ILC2s (g). (h,i) Flow cytometry analysis of KLRG1 by ILC2s (h) and quantification of the KLRG1 MFI (i). (j-m) Analysis of SI from 3-week-old *Vil1*^Cre^;*Il17rb*^fl/fl^ and *Il17rb*^fl/fl^ littermate mice, treated before and after birth with broad-spectrum antibiotics (MNVA, metronidazole, neomycin sulfate, vancomycin, and ampicillin) in drinking water. (j,k) Flow cytometry analysis of Ki-67 and Sm13 reporter expression by ILC2s (j) and quantification of IL-13 (Sm13)^+^ ILC2s (k). (l,m) Flow cytometry analysis of KLRG1 by ILC2s (l) and quantification of their KLRG1 MFI (m). (a,b) Data is representative of, and pooled from, 2 independent experiments. (c-e) Data pooled from 2 independent experiments. (f-m) Data is representative of, and pooled from 2 independent experiment. *, P 0.01– 0.05; **, P 0.01–0.001; ****, P < 0.0001.

### Tuft cell-intrinsic IL-17RB regulates tonic IL-25 bioavailability to prevent excessive ILC2 activation

Because tuft cells constitutively express *Il25* transcript, we hypothesized that deletion of the cognate receptor, IL-17RB, in tuft cells themselves results in increased IL-25 release and consequently the activation of ILC2s. To test if IL-25 is required for the elevated basal activation state of ILC2s in mice that lack IL-17RB in tuft cells, we generated *Vil1*^Cre^;*Il17rb*^fl/fl^;*Il25*^fl/fl^ mice. Indeed, absence of both IL-17RB and IL-25 abrogated the elevated expression of IL-13 and KLRG1 in 3-week-old mice (Fig. 6a-d). These results suggested a constant low-level production of IL-25 which is sensed by ILC2s. In agreement with prior data, we found that tuft cells readily express the *Il25*-tdTomato reporter also in young mice (Fig. 6e) [7]. To assess the consequence of IL-25 deficiency on the basal activation state of ILC2s, we analyzed *Vil1*^Cre^;*Il25*^fl/fl^ mice. Reduction in tdTomato expression in tuft cells of *Vil1*^Cre^;*Il25*^fl/fl^ mice revealed efficient Cre-mediated *Il25* deletion (Fig. 6e, Fig. S4a). When compared to ILC2s of their littermate controls, ILC2s from *Vil1*^Cre^;*Il25*^fl/fl^ mice showed a significant reduction in expression of IL-13, Ki-67 and KLRG1 (Fig. 6f-h), in accordance with our observations made in mice with ILC2-intrinsic *Il17rb*-deletion (Fig. 1). Furthermore, the frequency of tuft cells among epithelial cells was reduced when IL-25 was absent (Fig. 6i). Overall, this suggested that IL-25 is constitutively produced by tuft cells at low levels, which maintains basal ILC2 activation.

**Figure 6.**
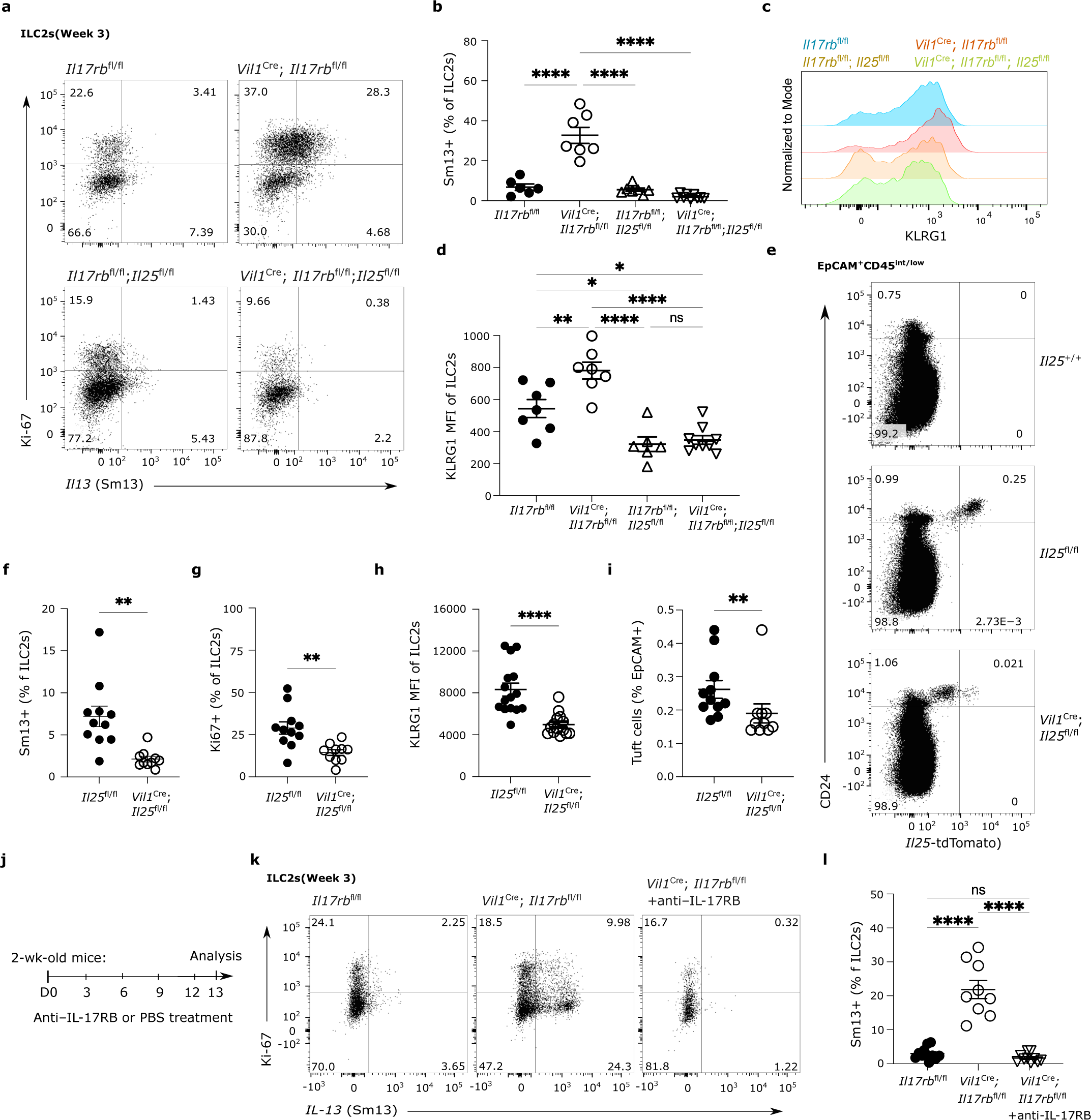
Tuft cell-intrinsic IL-17RB regulates tonic IL-25 bioavailability. (a-d) The small intestine (SI) was isolated from 3-week-old mice of the indicated genotypes and analyzed by flow cytometry. (a,b) Ki-67 and Sm13 reporter expression by ILC2s (a) and quantification of IL-13 (Sm13)^+^ ILC2s (b). (c,d) Analysis of KLRG1 expression by ILC2s (c) and quantification of their KLRG1 MFI (d). (e-i) Analysis of the SI from 3-week-old *Vil1*^Cre^;*Il25*^fl/fl^ and *Il25*^fl/fl^ littermate mice which also encode the *Il25*^tdTomato^ reporter (Flare25). (e) Expression of tdTomato and CD24 in CD45^int/low^EpCAM^+^ cells was determined by flow cytometry analysis and compared to that of a reporter-negative mouse. (f,g) Frequencies of IL-13 (Sm13)^+^ (f) and Ki-67^+^ (g) ILC2s. (h) Quantification of the KLRG1 MFI in ILC2s. (i) Quantification of tuft cells by flow cytometry. (j-l) Analysis of SI from young *Vil1*^Cre^;*Il17rb*^fl/fl^ and *Il17rb*^fl/fl^ littermate mice after five injections of PBS or anti-IL-17RB blocking antibody, according to the schematic in (j). Flow cytometry analysis of Ki-67 and Sm13 reporter expression by ILC2s (k) and quantification of IL-13 (Sm13)^+^ ILC2s (l). (a,d) Data pooled from 2 independent experiments. (e-i,k,l) Data is representative of, and pooled from, 3 independent experiments. *, P 0.01–0.05; **, P 0.01– 0.001; ****, P < 0.0001.

To further demonstrate that this tonic tuft cell IL-25 – ILC2 axis is causing an excessive activation in ILC2s when quenching by tuft cell IL-17RB is lacking, we treated 2-week-old *Vil1*^Cre^;*Il17rb*^fl/fl^ mice with anti-IL-17RB blocking antibody, enabling inhibition of IL-17RB engagement by IL-25 in ILC2s with temporal control (Fig. 6j). This treatment abolished the features of increased basal ILC2 activation in *Vil1*^Cre^;*Il17rb*^fl/fl^ mice (Fig. 6k,l). Because IL-17RB forms a receptor complex with the shared receptor component IL-17RA, absence of IL-17RB may result in altered receptor chain pairing of IL-17RA and a shift to enhanced signaling through the IL-17RA/IL-17RC complex, which has recently been suggested to promote secretory cell differentiation in the small intestine [29]. To exclude this possibility, we crossed *Vil1*^Cre^;*Il17rb*^fl/fl^ mice with *Il17rc*^−/−^ mice. These *Vil1*^Cre^;*Il17rb*^fl/fl^;*Il17rc*^−/−^ mice showed the same increase in IL-13 expression compared to *Vil1*^Cre^;*Il17rb*^fl/fl^ mice, demonstrating that signaling downstream of IL-17A and IL-17F is not involved (Fig. S4b,c). We concluded that tuft cells constitutively express *Il25* transcript and produce low amounts of IL-25 protein which contributes to basal ILC2 activation in the small intestine. Tuft cell IL-17RB expression regulates bioavailable IL-25 levels in the lamina propria, thereby influencing homeostatic regulation of the tuft cell–ILC2 circuit.

### Prolonged activation by IL-25 induces a hypoproliferative state in ILC2s

To better understand the functional consequences of uncontrolled tonic IL-25 release and the resulting prolonged stimulation of ILC2s, we treated adult *Vil1*^Cre^;*Il17rb*^fl/fl^ mice with succinate and assessed ILC2 activation 4 days later. IL-13 expression was similar between ILC2s from *Vil1*^Cre^;*Il17rb*^fl/fl^ and littermate control mice after stimulation of tuft cells with succinate, and it was comparable to the elevated expression observed in ILC2s from *Vil1*^Cre^;*Il17rb*^fl/fl^ mice at baseline (Fig. 7a). However, while both groups showed minimal baseline Ki-67 expression without treatment, ILC2s in *Vil1*^Cre^;*Il17rb*^fl/fl^ mice failed to upregulate Ki-67 upon succinate administration compared to their littermate controls which showed robust proliferative capacity (Fig. 7b). Because succinate induces tuft cell IL-25 release, we then bypassed tuft cells by directly injecting IL-25 into adult *Vil1*^Cre^;*Il17rb*^fl/fl^ and *Il17rb*^fl/fl^ control mice. Similar to the results with succinate, ILC2s from both genotypes robustly expressed IL-13, whereas the proliferative response, as assessed by Ki-67 expression, was reduced in ILC2s from *Vil1*^Cre^;*Il17rb*^fl/fl^ mice (Fig. 7c, d). We therefore concluded that ILC2s from *Vil1^Cre^*;*Il17rb*^fl/fl^ mice, which had been exposed to chronically elevated tonic IL-25 stimulation, became hyporesponsive towards additional IL-25 as elicited with tuft cell stimulation.

**Figure 7.**
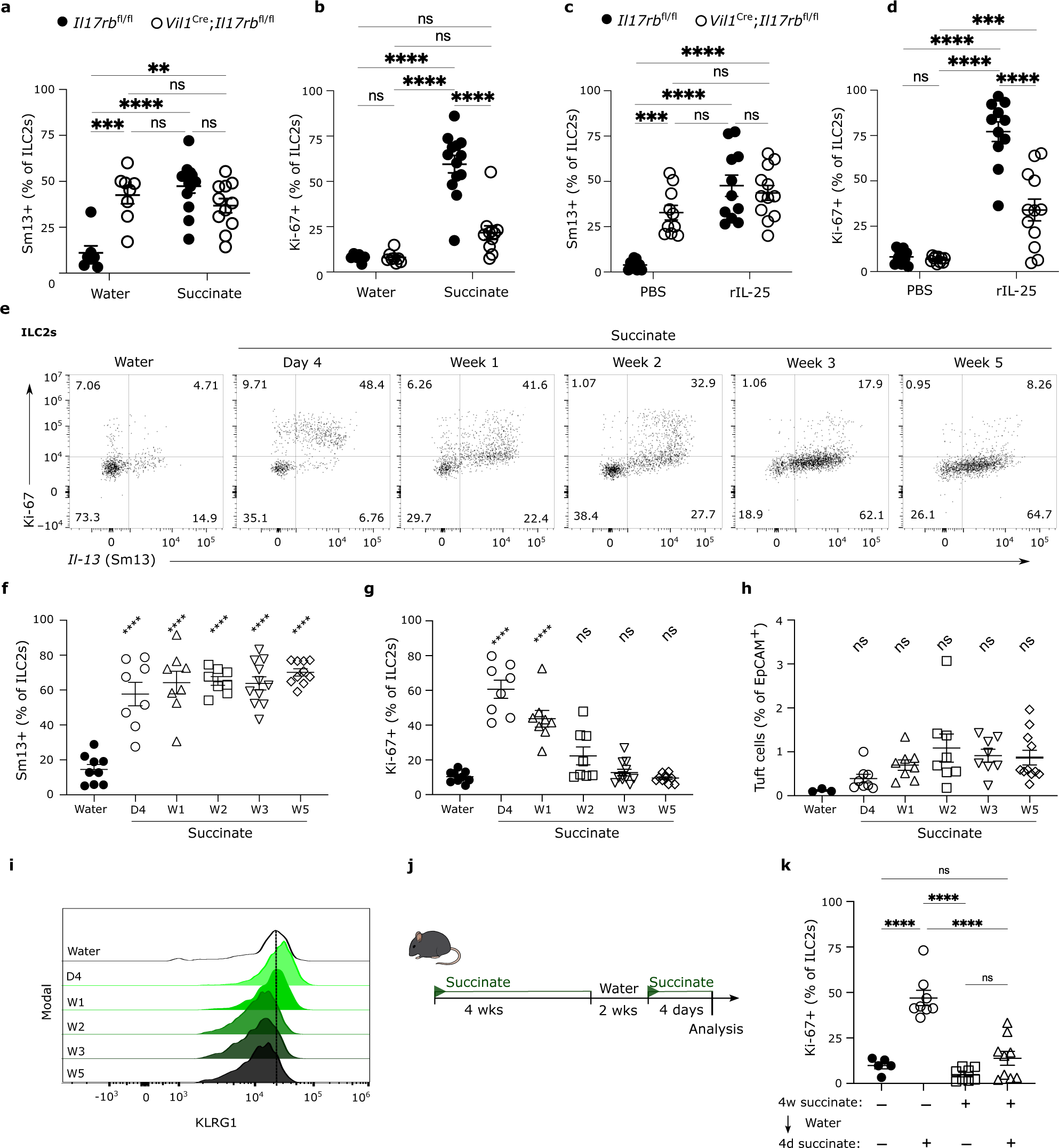
Prolonged activation by IL-25 induces a hypoproliferative state in ILC2s. (a-d) Adult *Vil1*^Cre^;*Il17rb*^fl/fl^ and *Il17rb*^fl/fl^ littermate mice were treated with succinate for 4 days (a,b) or rIL-25Fc (c,d). The percentage of IL-13 (Sm13)^+^ (a,c) and Ki-67^+^ (b,d) ILC2s in the small intestine (SI) was quantified by flow cytometry. (e-i) Wildtype mice encoding IL-13 (Sm13)-reporter were treated with succinate for the indicated amount of time (D, days; W, weeks) and cells from the SI were analyzed by flow cytometry. (e) Flow cytometry analysis of Ki-67 and IL-13 (Sm13)-reporter expression by ILC2s. (f,g) Frequencies of IL-13 (Sm13)^+^ (f) and Ki-67^+^ (g) ILC2s. (h) Percentage of DCLK1^+^CD24^+^ tuft cells among epithelial cells. (i) Representative histograms displaying KLRG1 expression by ILC2s. (j and k) Mice were treated with succinate for 4 weeks, followed by 2 weeks of regular drinking water, and another 4 days of succinate treatment, according to the schematic in (j). (k) Frequencies of Ki-67^+^ ILC2s in the small intestine lamina propria were analyzed by flow cytometry. (a-i) Data is representative of, and pooled from, 3 independent experiments. (k) Data pooled from 2 independent experiments. ** ≤ P = 0.01, *** ≤ P = 0.001, **** ≤ P = 0.0001.

To test this hypothesis in a model that did not depend on genetic deficiency in regulatory elements, we treated wildtype mice, that only encode reporter alleles, with succinate for a varying amount of time to promote and maintain tuft cell IL-25-mediated ILC2 activation (Fig. 7e). Regardless of the duration of stimulation, expression of IL-13 by ILC2s was maintained for as long as 5 weeks consistent with the persistent stimulation of tuft cells by succinate (Fig. 7e, f). In contrast, the percentage of Ki-67^+^ ILC2s exhibited a steady decrease following the initial peak, after 5 weeks reaching levels that resembled those found in ILC2s of untreated mice (Fig. 7e, g). This decline in Ki-67 expression occurred despite the subtle increase in tuft cell frequencies, and was paralleled by a reduction in ILC2 KLRG1 expression over time (Fig. 7h, i), overall recapitulating the findings observed under conditions of chronically elevated tonic IL-25 exposure caused by tuft cell *Il17rb*-deficiency in *Vil1*^Cre^;*Il17rb*^fl/fl^ mice.

Notably, the hypoproliferative state of ILC2s did not immediately revert. When wildtype mice were treated with succinate for 4 weeks, followed by regular drinking water for two weeks, before being re-exposed to succinate for 4 days, Ki-67 expression was still significantly lower compared to ILC2s from naïve mice that were acutely treated with succinate (Fig. 7j, k). A two-week duration is long enough for generation of new tuft cells, not previously exposed to succinate, making it unlikely that their response to the agonist would be atypical. We therefore concluded that prolonged stimulation with IL-25 is most likely responsible for inducing a state in resident ILC2s that hinders their sustained proliferative capacity.

### The hypoproliferative ILC2 state, triggered by suboptimal stimulation, is also associated with chronic helminth infection

ILC2s express receptors that let them integrate diverse molecular signals from their surrounding tissue environment [30]. We speculated that the stimulation of tuft cells with an agonist such as succinate, specifically triggering release of IL-25 but no robust production of other effector molecules that activate ILC2s, might be insufficient to enable proliferation over multiple cycles.

To test this hypothesis, we thought to provide stimulation with an additional signaling modality, in a highly specific manner, and to that end crossed YRS mice, expressing Cre in ILC2s, with a *Tg*^CAG-LSL-Gq-DREADD^ strain. This generated *Il5*^R/R^;*Tg*^Gq-DREADD^ mice in which ILC2s overexpressed a mutant hM3Dq G protein-coupled receptor (GPCR) that induces the canonical Gq pathway specifically following administration of the molecule clozapine-N-oxide (CNO) (Fig. 8a-c). This model enabled us to mimic GPCR–Gq–Ca^2+^–NFAT signaling as induced downstream of receptor activation by leukotrienes or neuromedin U (NMU) which are known to contribute to ILC2 activation [30]. *Il5*^R/R^;*Tg*^Gq-DREADD^ and *Il5*^R/R^ littermate control mice were then treated with succinate for 4 weeks to induce a hypoproliferative state, and CNO was injected twice over the last two days (Figure 8d). In agreement with our prior results, control mice without Gq-DREADD overexpression showed substantial IL-13 expression but lacked signs of proliferation as assessed by Ki-67 staining (Fig. 8e,f). In contrast, a majority of ILC2s from *Il5*^R/R^;*Tg*^Gq-DREADD^ mice were Ki-67^+^ (Fig. 8f). These results suggested that another, non-redundant, pathway together with IL-25 signaling is necessary for maintaining proliferative capacity.

**Figure 8.**
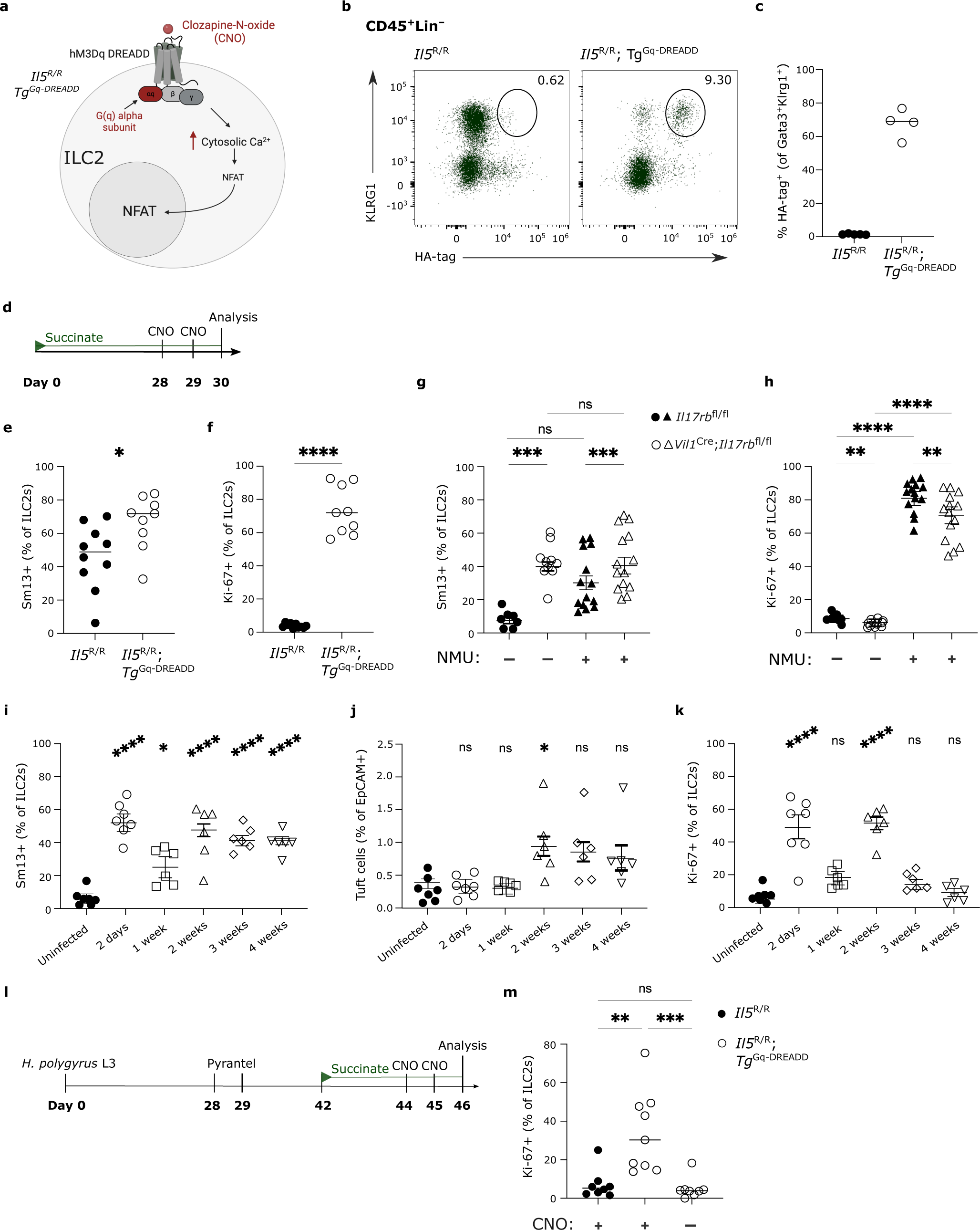
Stimulation of a non-redundant signaling pathway restores proliferative capacity in ILC2s. (a) Schematic of the signaling pathway engaged by the hM3Dq DREADD agonist CNO. (b and c) Small intestinal (SI) cells from *Il5*^R/R^ and *Il5*^R/R^;*Tg*^Gq-DREADD^ mice were analyzed by flow cytometry. (b) Expression of KLRG1 and hM3Dq DREADD (marked by HA tag) in ILC2s, gated on CD45^+^Lin^−^ cells. (c) Quantified percentage of hM3Dq DREADD^+^ ILC2s. (d) Schematic showing 30 day succinate treatment with CNO injections on two consecutive days prior to analysis; used in E and F. (e,f) Small intestinal (SI) cells from *Il5*^R/R^ and *Il5*^R/R^;*Tg*^Gq-DREADD^ mice were analyzed by flow cytometry and frequencies of IL-13 (Sm13)^+^ (e) and Ki-67^+^ (f) ILC2s quantified. (g, h) *Vil1*^Cre^;*Il17rb*^fl/fl^ and *Il17rb*^fl/fl^ littermate mice were treated with 20 μg NMU on two consecutive days and the percentages of IL-13 (Sm13)^+^ (g) and Ki-67^+^ (h) SI ILC2s were analyzed by flow cytometry one day later. (i–k) Mice were infected with *H. polygyrus* L3 and cells from the SI were analyzed on the indicated time points by flow cytometry. (i) Frequencies of IL-13 (Sm13)^+^ ILC2s. (j) Quantification of tuft cells by flow cytometry. (k) Frequencies of Ki-67^+^ ILC2s. (l, m) *Il5*^R/R^ and *Il5*^R/R^;*Tg*^Gq-DREADD^ mice were infected with *H. polygyrus* L3 and the worms were cleared by oral treatment with pyrantel pamoate, according to the schematic in (l). Two weeks later mice were treated with succinate in the drinking water for 4 days and injected with CNO on two consecutive days prior to analysis. Control groups consisted of *Il5*^R/R^ mice also treated with CNO, and *Il5*^R/R^;*Tg*^Gq-DREADD^ mice treated with PBS. (m) Frequencies of Ki-67^+^ ILC2s in the SI lamina propria were analyzed by flow cytometry. (a) Schematic made using biorender. (b,c,e,f, i-k, m) Data is representative of, and pooled from, 2 independent experiments. (g,h) Data is pooled from 3 independent experiments. * ≤ P = 0.05, ** ≤ P = 0.01, *** ≤ P = 0.001, **** ≤ P = 0.0001.

To test whether Ki-67 expression could, in a similar manner, also be restored in ILC2s from *Vil1*^Cre^;*Il17rb*^fl/fl^ mice, we treated adult animals with NMU, a time point in which their response to succinate or IL-25 was significantly impaired (Fig. 7b,d). NMU injection induced the expression of IL-13 and Ki-67 in ILC2 from control mice in accordance with prior studies (Fig. 8g,h) [31–33]. Although NMU did not further increase the already elevated baseline IL-13 expression in *Vil1*^Cre^;*Il17rb*^fl/fl^ mice, it potently increased the frequency of Ki-67^+^ ILC2s which were previously unable to proliferate when exposed to IL-25 or IL-25-inducing stimulus succinate (Fig. 8h). Together, these results demonstrate that the combined action of at least two non-redundant pathways are necessary to ensure full proliferative capacity in ILC2s.

Chronic helminth infection is marked by an elevation in tuft cell numbers, although this response alone proves ineffective in eradicating adapted parasites like *H. polygyrus* [3–5]. We hypothesized that a hypoproliferative state might also be induced in ILC2s under these conditions of persistent tuft cell– ILC2 stimulation, further contributing to the diminished type 2 responses, in addition to the direct immunomodulatory tactics employed by this parasite [34]. To assess this, we analyzed features of tuft cell–ILC2 circuit activation in YRS mice at different time points following infection with *H*. *polygyrus*. IL-13 expression by ILC2s was detected shortly after infection and persisted throughout the entire period when adult worms were present in the lumen, extending up to 4 weeks (Fig. 8i). The frequency of IL-13^+^ ILC2s decreased 1 week post infection, when worm larvae migrate away from the intestinal lumen and temporarily take up residence close to the outer muscularis layer, where they might be shielded from tuft cell detection (Fig. 8i). In parallel, tuft cell frequencies increased as reported previously (Fig. 8j). Notably, the percentages of Ki-67^+^ ILC2s mirrored the IL-13 expression pattern, characterized by an initial rise, a temporary decline, and a resurgence when adult worms colonize the lumen of the small intestine (Fig. 8k). However, the frequency of Ki-67^+^ ILC2s gradually decreased over the following two weeks, eventually reaching a level akin to that of uninfected mice (Fig. 8k).

To test whether ILC2s indeed acquired a lasting hypoproliferative state, we cleared the worms after 4 weeks and rested the mice for an additional two weeks, before treatment with succinate to stimulate IL-25-mediated ILC2 proliferation (Fig. 8l). By performing these experiments in *Il5*^R/R^ and *Il5*^R/R^;*Tg*^Gq-DREADD^ mice, we were also able to directly test the consequences of stimulating the Gq-DREADD pathway with CNO. Consistent with our prior observations, ILC2s from control groups failed to upregulate Ki-67, indicating an impaired proliferative response when stimulated via the succinate–IL-25 axis following helminth clearance (Fig. 8m). In contrast, injection of CNO during succinate stimulation significantly increased the frequency of Ki-67^+^ ILC2s in *Il5*^R/R^;*Tg*^Gq-DREADD^ mice (Fig. 8m). Overall, these results identify a hypoproliferative state in ILC2s following prolonged circuit activation due to chronic helminth infection, which could be overcome by providing stimulation via a separate, non-redundant, signaling pathway.

## Discussion

The investigation into the cytokine IL-25 has played a pivotal role in uncovering the existence of ILC2s and subsequently elucidating the involvement of tuft cells in initiating innate type 2 responses within the small intestine. Our research has yielded several crucial insights pertaining to the regulation of this cytokine in the predominant cell types orchestrating these responses. First, we identified key mechanistic steps governing the production/secretion of IL-25 by tuft cells, triggered by detection of succinate before subsequent activation of ILC2s. Second, building upon the implications from previous studies on IL-25 and ILC2s, we unequivocally demonstrated the ILC2-intrinsic requirement for IL-17RB during physiological stimulation of the tuft cell–ILC2 circuit. Third, we uncovered an unsuspected regulatory function of IL-17RB in tuft cells, controlling the levels of bioavailable IL-25 and thereby preventing excessive tonic stimulation of ILC2s resulting from constitutive expression of *Il25* transcript in tuft cells. Lastly, we described a hypoproliferative state in ILC2s induced by chronic IL-25 exposure, which could be offset by engagement of non-redundant pathways such as NFAT activation.

The discovery that tuft cells can respond to succinate, and subsequently stimulate small intestinal ILC2s in an IL-25-dependent manner, has provided the field with a straightforward and physiologically relevant model for studying the tuft cell–ILC2 circuit [7–9]. Tuft cells constitutively express *Il25 in vivo* and *in vitro*, even in the absence of agonistic molecules such as succinate [3]. In agreement with prior studies, possibly influenced by the presence of *Tritrichomonas* protists, we could confirm constitutive *Il25* expression, reported by a tdTomato knock-in allele. Tuft cells from neonatal mice housed in an SOPF facility free of *Tritrichomonas* expressed *Il25* as soon as they emerged in the small intestinal epithelium [7]. Constitutive expression of *Il25* transcript might reflect the anticipatory state in which tuft cells are engaged, marked also by concurrent expression of the enzymatic machinery for production of leukotrienes and acetylcholine [6, 8, 35, 36]. The constant *de novo* differentiation of tuft cells from the pool of LGR5^+^ stem cells, coupled with their short life span in the villus epithelium, likely requires such a state of preparedness to facilitate immediate and meaningful responses to luminal signals [28, 37]. Indeed, we found no evidence of *Il25* mRNA upregulation when tuft cells were stimulated with succinate. However, IL-25 protein becomes rapidly available to activate ILC2s, which we could prevent using an antibody that blocks the engagement of the receptor IL-17RB by IL-25.

The posttranscriptional regulation of IL-25 protein involves a series of intracellular events that are just beginning to be unravelled. Prior studies demonstrated the need for tuft cell PLCβ2 and TRPM5 – both highly expressed in tuft cells – in succinate-elicited, tuft cell-mediated, activation of ILC2s and the associated expansion of tuft cells [7–9, 38]. Our work highlights the critical role of Ca^2+^ release from intracellular stores in the ER, mediated by IP3R2, which we demonstrated using *Itpr2*^−/−^ mice and live cell calcium imaging of tuft cells in intact villi. The control IL-25 translation and/or its release by intracellular Ca^2+^ elevation, possibly also involving Ca^2+^-activated TRPM5-mediated depolarization, merit further investigation. Thus, unlike taste cells in the oral taste bud, which signal in an IP3R3-dependent manner, small intestinal tuft cells employ IP3R2 [24]. High expression of *Itpr2* in tuft cells from the human intestine indicates that this function might be conserved [26, 39]. In tracheal tuft cells, an elevation in intracellular calcium is linked to acetylcholine release that stimulates surrounding epithelial cells and triggers Ca^2+^ waves, thereby promoting ciliary activity and Cl^−^ secretion [22]. Small intestinal tuft cells may employ similar mechanisms to produce IL-25. Whether small intestinal tuft cells also depend on IP3R2 for TRPM5-mediated acetylcholine release and fluid secretion, or for helminth-induced leukotriene mobilization, requires further study [35, 36]. A recent report showing impaired helminth-evoked type 2 responses in mice lacking Lrmp, an ER-resident protein and possible interaction partner of IP3Rs, suggests that IP3R2 activation may indeed be a central signaling step downstream of multiple tuft cell stimuli [40].

Numerous reports have underscored the interdependence between tuft cells, IL-25, and ILC2s in facilitating type 2 responses, as observed in helminth infections, or during heightened luminal succinate levels [3–5, 7–9]. Early studies showed that IL-25 can activate ILC2s, also when isolated and stimulated *in vitro* [17, 18]. The high expression of IL-17RB in small intestinal ILC2s, coupled with numerous subsequent investigations employing IL-25 as a potent ILC2 agonist, has established the model whereby ILC2s directly respond to IL-25, known as the tuft cell–ILC2 circuit [41]. This assumption, however, has never undergone formal verification under conditions of physiological stimulation of the circuit *in vivo*. In this study, we utilized genetic tools enabling the specific deletion of IL-17RB in ILC2s. Our findings unequivocally demonstrate an intrinsic requirement for IL-17RB in ILC2s to foster proliferation and induce IL-13 expression upon succinate-mediated circuit stimulation. Although these observations were not tested in alternative settings of tuft cell stimulation, such as helminth infection, the presented data solidify ILC2s as a direct target of tuft cell-derived IL-25.

The presence of *Il17rb* transcript as an unexpected feature of tuft cells was noted already prior to the identification of their role as drivers of type 2 immunity, and later confirmed by transcriptional profiling of tuft cells in mice and humans [8, 26, 39, 42, 43]. Our results provide further evidence for an anticipatory state of tuft cells wherein they sustain *Il25* transcripts in the absence of luminal triggers, yet in which intrinsic IL-17RB expression regulates tonic levels of bioavailable IL-25. The feed-forward nature of the circuit likely requires tight control to enable a rapid response while still preventing inappropriate steady-state stimulation, which is further supported by mechanisms constraining activation in ILC2s such as A20 and CISH [7, 19]. This threshold can be overcome by tuft cell agonists like succinate that trigger swift IL-25-mediated circuit activation. Whether agonistic signaling lowers activity of the regulatory breaks, or simply stimulates the production of IL-25 protein to levels exceeding the quenching activities, requires further investigation. Our studies also did not address whether tuft cell IL-17RB only sequesters IL-25 or if this interaction governs additional regulatory signaling in tuft cells. Recent studies linking IL-25 and IL-17RB with intestinal cancer warrant further characterization of this cytokine axis [44–46].

Intriguingly, we also describe a hypoproliferative state in ILC2s resulting from continuous stimulation by IL-25, induced either by genetic deletion of IL-17RB in tuft cells, or by prolonged succinate exposure. This decline in ILC2 proliferation may reflect natural adaptation to states of prolonged tuft cell stimulation, adding an additional layer of control to prevent excessive circuit activity. Indeed, continuous low-grade stimulation of tuft cells by succinate or possibly other microbial agonists may be more common in a natural environment with dynamic alterations in the luminal state compared to the stable and low type 2 tone in mice housed in barrier facilities. Such circuit activation might be sufficient to provoke responses mediating adaptive alteration in antimicrobial programs [47, 48], yet may not promote the very strong expansion of tuft and goblet cells seen with *N. brasiliensis* infection [3–5]. We hypothesize that the ILC2 response is controlled by the sum of activating signaling pathways engaged, and that they may integrate these during early activation in a way that subsequently governs the magnitude of proliferation, similar to what was proposed for T cells [49]. Prolonged suboptimal stimulation by IL-25 resulted in impaired proliferation when acutely challenged with succinate later on, suggesting that ILC2s record conditions of prior stimulation for a certain period. This may involve epigenetic alterations and possibly reflect adaptation to repeated colonization or bursts in succinate-producing microbes, not requiring strong ILC2 expansion. Such reduced sensitivity resembles the law of ‘initial value’ by J. Wildner *et al.*, conceptualizing how the basal state of activity can determine if a subsequent stimulus is able to elicit a biological response, recently discussed in the context of T cell responses [50, 51]. Notably, although prior studies reported states of ILC2 exhaustion, our results with engaging synergistic NFAT signaling pathways show a rapid reversal of the hypoproliferative state when an optimal amount of ILC2 stimulation is available [52, 53].

A consequence of ILC2s transitioning to a hypoproliferative state may be the impaired clearance of adapted helminths such as *H. polygyrus*. Under these circumstances, suboptimal levels of ILC2 stimuli may persist despite the continued abundance of luminal worms. Consequently, the induction of hypoproliferation could represent an additional manipulation in host cell responsiveness, akin to the recently described shaping of the epithelial response to IL-13 [54]. Notably, the antagonism of IL-33 activity may contribute to the suboptimal level of ILC2 stimulation, facilitating the hypoproliferative state. Further studies are needed to dissect the alterations induced by chronic helminth infections in the activity of individual pathways acting on ILC2s, including those involving epithelial cytokines, eicosanoids, and neuropeptides [30]. These mechanisms may unveil the elegant immunomodulatory potential of helminths, beyond the direct antagonistic action of secreted parasite products [34]. Overall, such adaptive responses in the host may prove beneficial if productive clearance cannot be achieved, and excessive chronic type 2 stimulation might lead to undesirable effects.

## Methods

### Resource sharing

Further information regarding resources/reagents, or sharing thereof, should be directed to the corresponding author, Christoph Schneider.

### Mice

We used the following mouse strains: *Vil1*^Cre^ (B6.Cg-Tg(Vil1-cre)997Gum/J; JAX, 004586); *Nmur1*^iCre-eGFP^ ([21], from D. Artis and C. Klose); *Il17rb*^fl/fl^ ([15], from U. Siebenlist); *Il17rb*^−/−^ mice with global *Il17rb* deletion were generated from *Vil1*^Cre^; *Il17rb*^fl/fl^ mice in which Cre is active with high frequency in the male germline; *Itpr2*^−/−^ ([55], from A. Saab); *Trpm5*^Cre^;*R26*^GCaMP6f^ [22, 23]; *Il17rc*^−/−^ (from S. Leibundgut-Landmann); *Il5*^Red5^ (B6(C)-*Il5^tm1.1(iCre)Lky^*/J; JAX, 030926); *Il25*^fl-tdTomato^ ([3], B6(C)-*Il25^tm1.1Lky^*/J); *Il25*^iCre^ ([28], B6(C)-*Il25^tm2.1(cre)Lky^*/J); *Tg*^CAG-lsl-Gq-DREADD^ (B6N;129-Tg(CAG-CHRM3*,-mCitrine)1Ute/J; JAX, 026220). Except for *Trpm5*^Cre^;*R26*^GcaMP6f^ mice [22, 23], all lines were crossed to *Arg1*^Yarg^;*Il13*^Smart13^ double reporter (B6.129S4-*Arg1^tm1Lky^*/J; JAX, 015857; B6.129S4(C)-*Il13^tm2.1Lky^*/J; JAX, 031367). All mice were on a C57BL/6 background. Mice were bred and housed at the University of Zurich, Laboratory Animal Sciences Center in Zurich, Switzerland, under specific pathogen-free conditions, free of *Tritrichomonas*. Animals were housed in individually ventilated cage units containing autoclaved bedding and nesting material. Animal experiments were reviewed and approved by the cantonal veterinary office of Zurich (permit numbers 054/2022 and 111/2022), and in accordance with the guidelines established by the German Animal Welfare Act, European Communities Council Directive 2010/63/EU, the institutional ethical and animal welfare guidelines of Saarland University (approval number of the Institutional Animal Care and Use Committee: CIPMM-2.2.4.1.1). Age-matched mice of both sexes were at the age of 6 weeks or older, except for the indicated experiments with pups. Experiments were performed with mice randomly allocated to groups and without investigator blinding. No statistical methods were used to predetermine sample sizes (sample sizes similar to those reported in previous publications). No data points or animals were excluded from the analyses for any reason other than an animal becoming sick, injured, or dying for reasons unrelated to the experiment.

### Mouse treatments

**Succinate treatment:** 100 mM succinic acid (S3674, Sigma) was dissolved in autoclaved water, pH neutralized with NaOH, and filtered (0.22 μm). Mice were treated for the indicated duration with succinate instead of regular drinking water, and the solution was exchanged weekly. **Anti-IL-17RB treatment:** Monoclonal blocking antibody against IL-17RB (Clone D9.2) [18] was purified in house and administered i.p. at the indicated time points (antibody injection dose: adult mice, 200 μg; pups, 50 μg). **Antibiotic treatment:** Breeders received drinking water supplemented with ampicillin (1 mg/mL), metronidazole (1 mg/mL), neomycin (1 mg/mL), vancomycin (0.5 mg/mL), and sucrose (1 %) starting approximately 3 weeks prior to the birth of a litter until offspring reached 3 weeks of age. **IL-25 treatment:** Adult mice received 1 μg recombinant mIL-25–hFc fusion protein (purified in house) i.p. on two consecutive days prior to euthanasia. **CNO treatment:** Adult mice received i.p. injections of water-soluble Clozapine N-oxide (CNO) dihydrochloride, 20 μg in 200 μL, 2 consecutive days before euthanasia. **NMU treatment:** Adult mice received i.p. injections of mouse neuromedin U 23 peptide (NMU) (H-FKAEYQSPSVGQSKGYFLFRPRN-NH2, Mimotopes, Australia), 20 μg in 200 μL, 2 consecutive days prior to euthanasia. ***H. polygyrus* infection**: Adult mice were infected with 200 infectious *H. polygyrus* L3, in 200 μL, by oral gavage. For worm clearance, mice were treated on 2 consecutive days by oral gavage with 2 mg pyrantel pamoate (Perrigo, 36200).

### Tissue processing

Adult mice were euthanized through carbon dioxide, and pups by decapitation. **Small intestine:** As previously described [56], single cell suspensions were prepared using either the “basic protocol”, or the “alternate protocol 1” depending on expected inflammatory status (*i.e.* resting mice = basic protocol, following type 2 stimulation = alternate protocol). It proved important for successful staining of IL-17RB that the basic protocol was used. In short, duodenal tissue from adult mice, or complete intestine from pups, was opened longitudinally and incubated for 15 minutes in Ca^2+^/Mg^2+^ -free HBSS buffer supplemented with FCS (2 %), HEPES (10 mM) and DTT (5 mM). Tissues were then transferred into a fresh Ca^2+^/Mg^2+^ -free HBSS solution supplemented with FCS (2 %), HEPES (10 mM), EDTA (5 mM), and incubated for another 15 minutes prior to vortexing. The supernatant containing epithelial cells, was subsequently filtered (100 μm) into cold FACS buffer. The last step was repeated for a total incubation time in EDTA-buffer of 30 minutes. The epithelial fraction was kept on ice from this point on. Tissue samples were moved to a Ca^2+^/Mg^2+^ -containing HBSS solution supplemented with FCS (2 %), HEPES (10 mM), and incubated for another 10 minutes. Tissues were subsequently placed in Ca^2+^/Mg^2+^ -containing HBSS media supplemented with FCS (2 %), HEPES (10 mM), Liberase TM (100 μg/mL) and DNase 1 (30 μg/mL), where they were manually cut into small pieces. Following a 20-minute incubation, mechanical disassociation using GentleMACS C tubes (Miltenyi Biotec) and program m_intestine_01 on the gentleMACS dissociator (Miltenyi Biotec) was employed. Samples were then filtered (100 μm) and kept on ice for subsequent staining. All incubations were performed at 37 °C under gentle agitation.

### Flow cytometry and cell sorting

Single cell suspensions were incubated with Fc-receptor blocking antibody diluted in FACS buffer (1x PBS, supplemented with 5 % FCS and 0.1 % sodium azide). They were subsequently resuspended, for 20 minutes (particularly important for staining of tuft cell IL-17RB), in a cocktail of antibodies directed towards surface antigens. Lineage markers included: CD11c, CD19, NK1.1, Ter119, CD49b (DX5). See “resource table” for further information on antibodies used. Cells were then incubated with either DAPI or a zombie red dye to enable later exclusion of dead cells. In the event of live cell acquisition, samples were here resuspended in FACS buffer and immediately run on one of the three machines mentioned below. In case of continued intracellular staining, epithelial samples were fixed for 3 minutes using PFA (4 %), and lamina propria suspensions fixed for 10 minutes using the Foxp3/Transcription factor staining set (Thermo Fisher 00-5523-00). Cells were then resuspended in 1x permeabilization buffer (Thermo Fisher 00-8333-56) containing all antibodies directed towards intracellular targets. Cells were kept on ice throughout the staining procedure. Acquisition was done using either the spectral analyzer Cytek Aurora, the BD FACSymphony, or (for cell sorting) the FACSymphony S6. During sorting, roughly 10K ILC2s (defined as CD45^+^, lineage^−^ (here including also CD11b, CD8a and SiglecF), CD3^−^, CD4^−^, Thy1.2^+^ and KLRG1^+^) were sorted into RLT Lysis buffer (Qiagen) and pooled (2x) prior to RNA extraction.

### Immunofluorescence

The small intestinal tissue was flushed with PBS, cut open longitudinally, and fixed for 2+ hours in paraformaldehyde (4 % w/v) at 4 °C. This was followed by a PBS wash and an O/N incubation in sucrose (30 % w/v) at 4 °C. Folded into “swiss-rolls”, tissues were subsequently embedded in OCT and stored at –80 °C before sectioning (6 μm) on a Leica CM1850 Cryostat (Leica Biosystems). Sections were incubated for 1 h, RT, in blocking buffer (1x PBS supplemented with 2 % BSA, 0.1 % Triton x100, and 5 % goat serum). All staining was subsequently performed in blocking buffer. Incubation with primary antibodies was at 4 °C O/N, and incubation with secondary antibodies for 1 h at RT. Please see “resource table” for further information on antibodies used. DAPI was added separately for 5 minutes and samples were washed with PBS. For mounting, Vectashield mounting media was diluted 1:3 in a glycerol solution with added Tris (50 mM). Images were acquired on a fluoresence microscope (20x Leica DMi8; Leica Camera AG). Fiji (Image J) software was used to acquire and process the images.

### RNA sequencing

RNA was isolated from FACSorted samples using the Quick-RNA Microprep kit (Zymo Research; R1050). For RNA sequencing, samples were then stored for less than one week in –80 °C before being shipped on dry ice to Novogene (Germany). Samples underwent SMARTer amplification, prior to sequencing on an Illumina platform, where paired end reads were generated. We estimated transcript counts using Kallisto v0.48.0 [57] with mm10 reference genome and used tximport v1.28.0 [58] to import them in R v4.3.2. Based on normalized counts we removed 3 outlier samples using *PcaProj* function from rrcov v1.7.4 [59] R package and proceeded with differential expression analysis using DESeq2 v1.40.1 [60]. Normalized counts from DESeq2 for the selected genes were used to generate heatmaps with ComplexHeatmap v2.16.0 [61].

### Confocal Ca^2+^ imaging of tuft cells in an en-face ileum and scraped villi preparation

Mice (6 - 22 weeks old, both sexes) were decapitated after CO_2_ anesthesia. The intestine was excised and cleansed by perfusing with an extracellular solution containing (in mM): 136.5 NaCl, 5.6 KCl, 2.2 CaCl_2_, 1 MgCl_2_, 10 HEPES adjusted to pH 7.4 (NaOH) and 290 mOsm (∼ 10 mM glucose). The ileum, approximately the last quarter of the intestine, was divided into 4 sections and placed on small strips of tissue paper (Kimtech Science) to prevent the en-face ileum from curling after cutting the tube-like structure open. The 4 en-face ileum parts were then placed into separate recording chambers (Luigs & Neumann, Germany) and secured by a tissue slice holder (harp) to prevent movement caused by the speed of the solution flow through the chamber [62, 63]. In case of the scraped villi preparation, the whole ileum was scraped using a scalpel blade no. 10. The scraped villi were directly added to a solution containing the Ca^2+^ indicator, slightly triturated and placed into the recording chamber containing an acid-washed glass coverslip coated twice with 0.01 % poly-L-lysine (Sigma P-6282; CAS # 25988-63-0). The scraped villi were also secured by a tissue slice holder (harp) to prevent movement.

Intracellular Ca^2+^ was monitored with either GCaMP6f expressed in the tuft cells of *Trpm5*^Cre^;*R26*^GcaMP6f^ mice or the Ca^2+^ indicator Cal630-AM loaded into tracheal epithelial cells using a similar technique as described previously for the tracheal epithelium [22]. Cal630-AM (AAT-Bioquest CAT# 20720) was dissolved in a solution of DMSO and freshly prepared 20% Pluronic F-127 in DMSO, then further diluted in extracellular solution (see above), and briefly sonicated. The en-face ileum or scraped villi were subsequently incubated in the Cal630-AM loading solution having a final concentration of 4.1 μM Cal630-AM, 0.15 % DMSO and 0.01 % Pluronic F127 for 90-120 min at room temperature. Imaging experiments were performed using an upright confocal laser scanning microscope (Zeiss LSM 880 INDIMO) with a Plan-Apochromat 20X/1.0 water immersion objective. Excitation wavelength for GCaMP6 was 488 nm and emitted fluorescence was collected between 500 and 540 nm. For calcium imaging in the IP3R2 mice, the tuft cells were identified using an excitation wavelength for tdTomato being 561 nm and using emission wavelength between 565 - 593 nm. The excitation wavelength for the Ca^2+^ indicator Cal630 was 594 nm and emitted fluorescence was collected between 600 - 690 nm. The excitation wavelength used for the Ca^2+^ indicator Cal630 is unable to excite the tdTomato fluorescence. To prevent Cal630 emissions from being collected due to a possibility of exciting the indicator by the 561 nm laser, the emission wavelength for tdTomato was limited to a maximum of 594 nm. All scanning head settings were kept constant during each experiment. Optical sections were 12.1 μm thick and were kept constant in all recordings. Images (512 x 512 pixels/frame) were acquired every 0.631 s. The following criteria for stimulus-induced Ca^2+^ responses were applied: (1) A response was defined as a stimulus-dependent deviation of either GCaMP6f or Cal630 fluorescence signal that exceeded twice the SD of the mean of the baseline fluorescence noise. (2) A response had to occur within 2 min after stimulus application. In time series experiments, ligand application was repeated to confirm the repeatability of a given Ca^2+^ response. GCaMP6f or Cal630 fluorescence changes of individual cells are expressed as relative fluorescence changes, i.e. ΔF/F (F being the average during control stimulation with extracellular solution). The maximum change in relative fluorescence is indicated as Fpeak. Images were processed with ZenBlack software (Zeiss) and analyzed using Fiji/ImageJ (NIH), Igor Pro (Wavemetrics) and Originlab (Origin) software. Through the Igor Pro software package, user-defined functions in combination with an iterative Levenberg-Marquardt nonlinear, least-squares fitting routine were applied to the data. Dose-response curves were fitted by the equation:

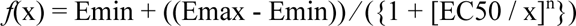

where x is the drug concentration, Emin the baseline response, Emax the maximal response at saturating concentrations, and EC50 the drug concentration that produces 50 % of the maximal response with slope n being the Hill coefficient of the sigmoid curve.

The intestinal epithelium was stimulated successively using bath application. Chemostimuli for Ca^2+^ imaging were prepared fresh daily and diluted in extracellular solution. The final succinate (CAS# 6106-21-4) concentrations were 0.3, 1.0, 3.0, and 10 mM. The impact of intracellular Ca^2+^ stores was examined with CPA (cyclopiazonic acid, 10 μM, CAS# 18172-33-3). Final DMSO concentrations (< 0.1%, vol/vol) were tested in control solutions and had no effects. All chemicals were obtained from Merck (previously Sigma/Aldrich) if not otherwise stated. The extracellular 60 mM KCl solution contained (in mM): 82.1 NaCl, 60 KCl, 2.2 CaCl_2_, 1 MgCl_2_, 10 HEPES adjusted to pH 7.4 (NaOH) and 290 mOsm (∼ 10 mM glucose).

### Statistical analysis

Unless otherwise indicated, data is pooled from 2 or more repeat experiments and displayed as the mean values (+/– SEM) in all graphs, with n’s reflecting biological replicates. For statistical analysis of data generated from 2 groups, a two-tailed (unpaired, unless otherwise indicated in figure legend) Student’s *t* test was used when normal distribution could be assumed, and a Mann-Whitney U test used in cases where it could not. For statistical analysis of more than 2 groups, an ordinary one-way ANOVA with Tukey’s multiple-comparison test was used throughout, apart from a two-way ANOVA being used for data containing two independent variables (see figure 3, figure legend). Results were deemed significant if P<0.05. Except in the case of major experimental error, no data was excluded from analysis. Intragroup variation was not assessed. All statistical analysis, with exception for data generated in RNA sequencing, was performed using Prism 10 (GraphPad Software) or Origin Pro (OriginLab Corporation, Northampton, MA, USA).

### Resource table

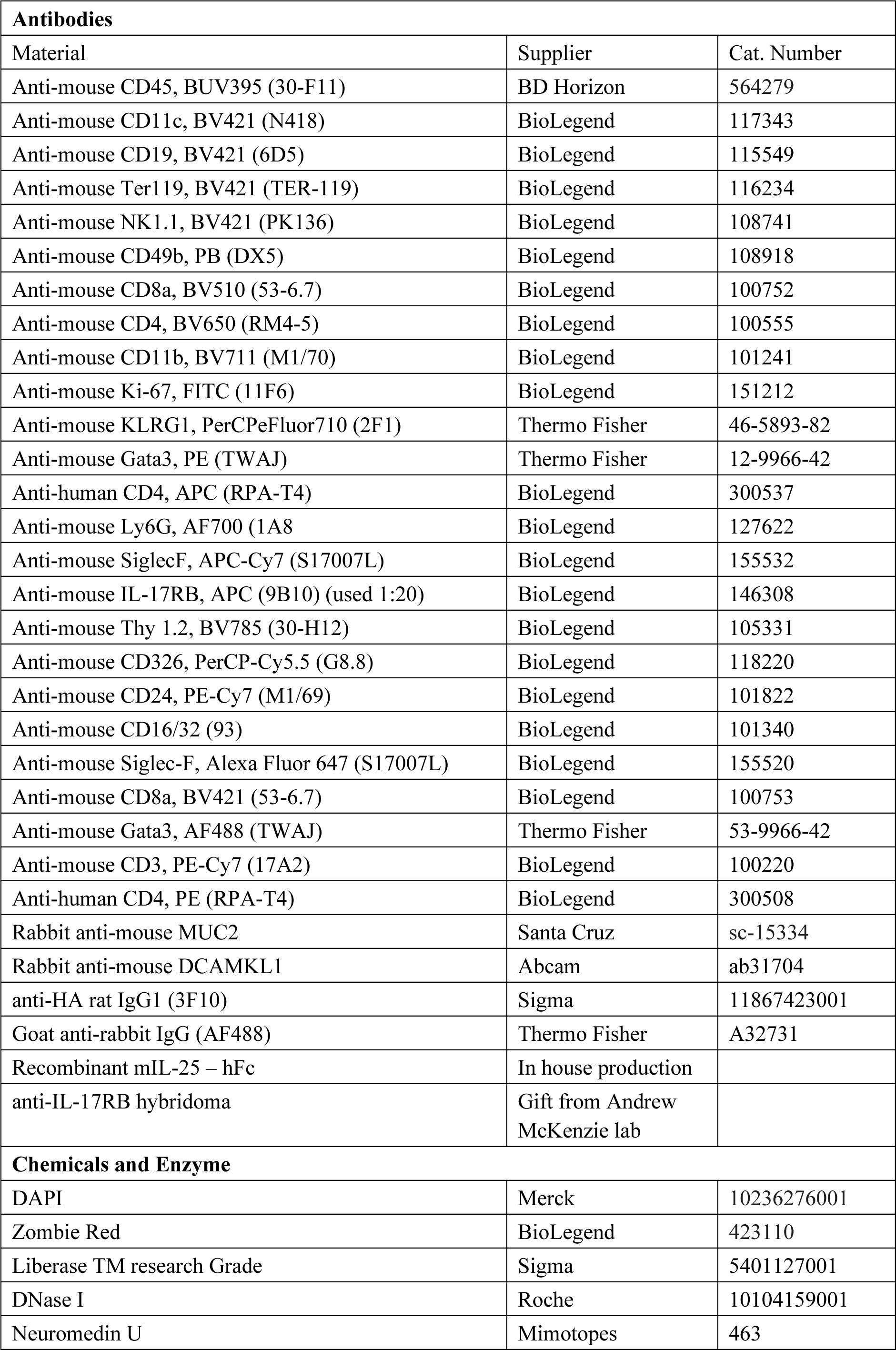
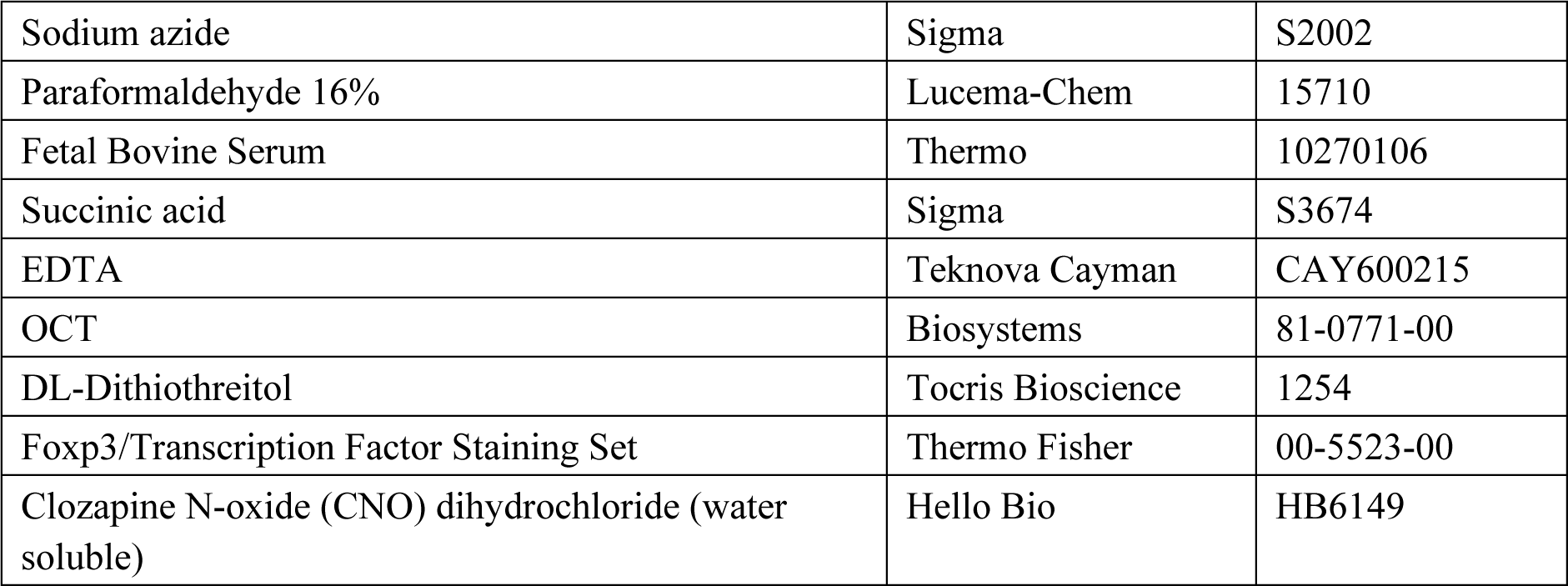

## Acknowledgements

We thank all members of the Schneider lab and Manfred Kopf for helpful discussion and input on this manuscript; Brigitte Herzog, Reto Kroeschell, Luca Plan, Lubor Borsig, and Thierry Hennet for technical expertise and access to equipment; Andrew McKenzie for providing the anti-IL-17RB hybridoma; Salomé LeibundGut-Landmann, Richard Locksley, Aiman Saab, and Uli Siebenlist for providing mice.

This work was supported by grants from the Swiss National Science Foundation (Eccellenza grant 194216), the Peter Hans Hofschneider Professorship for Molecular Medicine, the Foundation for Research in Science and the Humanities at the University of Zurich, and the Olga Mayenfisch Foundation to C.S.; the Deutsche Forschungsgemeinschaft (DFG) Grant Sonderforschungsbereich-Transregio TRR 152 to U.B., T.L.-Z., and F.Z.; the DFG-Instrumentation Grant INST 256/427-1 FUGB to T.L.-Z.; the European Research Council Starting Grant (ERCEA; 803087) and the German Research Foundation (DFG; FOR2599 Project-ID 22359157, CRC/TRR 241 Project-ID 375876048, SPP1937 - KL 2963/3-1 and KL 2963/2-1) to C.S.N.K.; the Jill Roberts Institute for Research in IBD, Kenneth Rainin Foundation, the Sanders Family Foundation, Rosanne H. Silbermann Foundation, CURE for IBD, the Allen Discovery Center program, a Paul G. Allen Frontiers Group advised program of the Paul G. Allen Family Foundation, and the US National Institutes of Health (DK126871, AI151599, AI095466, AI095608, AR070116, AI172027, DK132244) to D.A.. T.A. was supported by a UZH Candoc Grant. J.L.M. was supported by the «Personenfoerderung» Program of the Department of Surgery at the University Hospital Basel.

## Author contributions

X.F. and T.A. conceived the study, designed, and performed experiments, analyzed data and wrote the manuscript with C.S., J.G., P.F., J.L.M., D.C., A.L., J.D.T. and N.B. performed experiments. I.B. helped with RNA-seq data analysis. T.L.-Z and F.Z. designed and performed experiments, and provided access to critical tools. U.B., D.A., and C.S.N.K. provided critical tools. C.S. directed the study and wrote the paper with X.F. and T.A..

