## Supplementary figures for "Tuft cell IL-17RB restrains IL-25 bioavailability and reveals context-dependent ILC2 hypoproliferation"

### Supplementary Figure Legends

**Supplementary figure 1. Impaired succinate-mediated ILC2s activation in *Il5*$^{\mathrm{R/+}}$;*Il17rb*$^{\mathrm{fl/fl}}$ mice.** (a) Flow cytometry gating of small intestinal ILC2s, throughout the paper defined as either Gata3$^{+}$KLRG1$^{+}$ (fixed panel, filled line), or KLRG1$^{+}$ (live panel, dashed line). (b,c) The small intestine (SI) of naïve *Il5*$^{\mathrm{R/+}}$;*Il17rb*$^{\mathrm{fl/+}}$, *Il5*$^{\mathrm{R/+}}$;*Il17rb*$^{\mathrm{fl/fl}}$ and *Il17rb*$^{-/-}$ mice was analyzed. (b) Red5 or IL-17RB expression by CD3$^{+}$CD4$^{+}$ cells, gated as in A . (c) Expression of KLRG1 and *Arg1*$^{\mathrm{YFP}}$, IL-5 (Red5), and IL-13 (Sm13) reporters by Lin$^{-}$ cells. (d) *Il5*$^{\mathrm{R/+}}$;*Il17rb*$^{\mathrm{fl/+}}$, *Il5*$^{\mathrm{R/+}}$;*Il17rb*$^{\mathrm{fl/fl}}$ and *Il17rb*$^{-/-}$ mice were treated with succinate for 4 days and flow cytometry plots illustrating ILC2 gating and their expression of Ki-67 and IL-13 (Sm13) are shown. (a-c) Data is representative of 2 independent experiments. (d) Data is representative of 4 independent experiments. *, P 0.01–0.05; **, P 0.01–0.001; ****, P < 0.0001.

**Supplementary figure 2. Visualization of *Il25* expression and Ca$^{2+}$ signaling in tuft cells.** (a) Flow cytometry gating of small intestinal tuft cells, throughout the paper defined as either DCLK1$^{+}$CD24$^{+}$ (fixed panel, filled line), or *Il25*$^{\mathrm{tdTomato+}}$CD24$^{+}$ double positive (live panel, dashed line). (b,c) Tuft cell Ca$^{2+}$ activity of the ileum was imaged with an *ex vivo* whole mount (en-face view of the ileum) preparation from *Trpm5*$^{\mathrm{Cre}}$;*R26*$^{\mathrm{GCaMP6f}}$ mice. Tuft cells (*green*) can be identified based on *Trpm5* promoter-dependent expression of GCaMP6f (c, *arrows*). (d) Tuft cell Ca$^{2+}$ activity of the ileum was imaged with an *ex vivo* whole mount preparation from *Il25*$^{\mathrm{tdTomato}}$ (Flare25) mice using a Cal630 Ca$^{2+}$ indicator dye. Tuft cells (white box) can be identified based on their distinct *Il25* (*red*) and CD24 (*green*) expression of GCaMP6f (b, *arrows*). All experiments have been repeated a minimum of two times.

**Supplementary figure 3. ILC2s with an activated phenotype in mice with tuft cells *Il17rb*-deficiency.** (a,b) The small intestine (SI) of naïve *Vil1*$^{\mathrm{Cre}}$;*Il17rb*$^{\mathrm{fl/fl}}$ and *Il17rb*$^{\mathrm{fl/fl}}$ littermate mice was analyzed. Expression of IL-13 (Sm13) and *Arg1*$^{\mathrm{YFP}}$ reporters (a) or IL-17RB (b) by CD45$^{+}$Lin$^{-}$ lamina propria cells. (c) Frequencies of IL-17RB$^{+}$ tuft cells were analyzed in mice of the indicated genotypes by flow cytometry. (d) Tuft cell percentage in the epithelium of SI from naïve *Il25*$^{\mathrm{iCre/+}}$;*Il17rb*$^{\mathrm{fl/fl}}$ and *Il17rb*$^{\mathrm{fl/fl}}$ littermate mice was determined by flow cytometry. (a,b) Data is representative of 2 independent experiments. (c) Displayed data is from one experiment, representative of 2 independent experiments. (d) Data is pooled from 2 independent experiments. *, P 0.01–0.05; **, P 0.01–0.001; ****, P < 0.0001.

**Supplementary figure 4. Elevated tonic ILC2 stimulation in *Vil1*$^{\mathrm{Cre}}$;*Il17rb*$^{\mathrm{fl/fl}}$ mice is independent of *Il17rc*.** (a) Analysis of the small intestine (SI) from 3-week-old *Vil1*$^{\mathrm{Cre}}$;*Il25*$^{\mathrm{fl/fl}}$ and *Il25*$^{\mathrm{fl/fl}}$ littermate mice which also encode the *Il25*$^{\mathrm{tdTomato}}$ reporter (Flare25). tdTomato MFI of EpCAM$^{+}$ CD24$^{+}$SiglecF$^{+}$ tuft cells were determined by flow cytometry and compared to that in a reporter-negative mouse. (b,c) The SI was isolated from 3-week-old mice of the indicated genotypes and analyzed by flow cytometry.

Enumeration of IL-13 (Sm13)$^{+}$ ILC2 percentage (b) and quantification of the KLRG1 MFI in ILC2s (c). (a) Data pooled from 3 independent experiments. (b,c) Data pooled from 2 independent experiments. *, P 0.01–0.05; **, P 0.01–0.001; ****, P < 0.0001.

**Supplementary movie 1. Succinate-induced Ca2+-response in tuft cells.** Example of succinate-induced intercellular $Ca^{2+}$ responses in *Trpm5*$^{Cre}$;*R26*$^{GCaMP6f}$ tuft cells of the ileum from an *ex vivo* whole mount (en-face view of the ileum) preparation from *Trpm5*$^{Cre}$;*R26*$^{GCaMP6f}$ mice. The speed of the movie was increased by 10x. Data is representative of multiple independent experiments.

Figure 1S

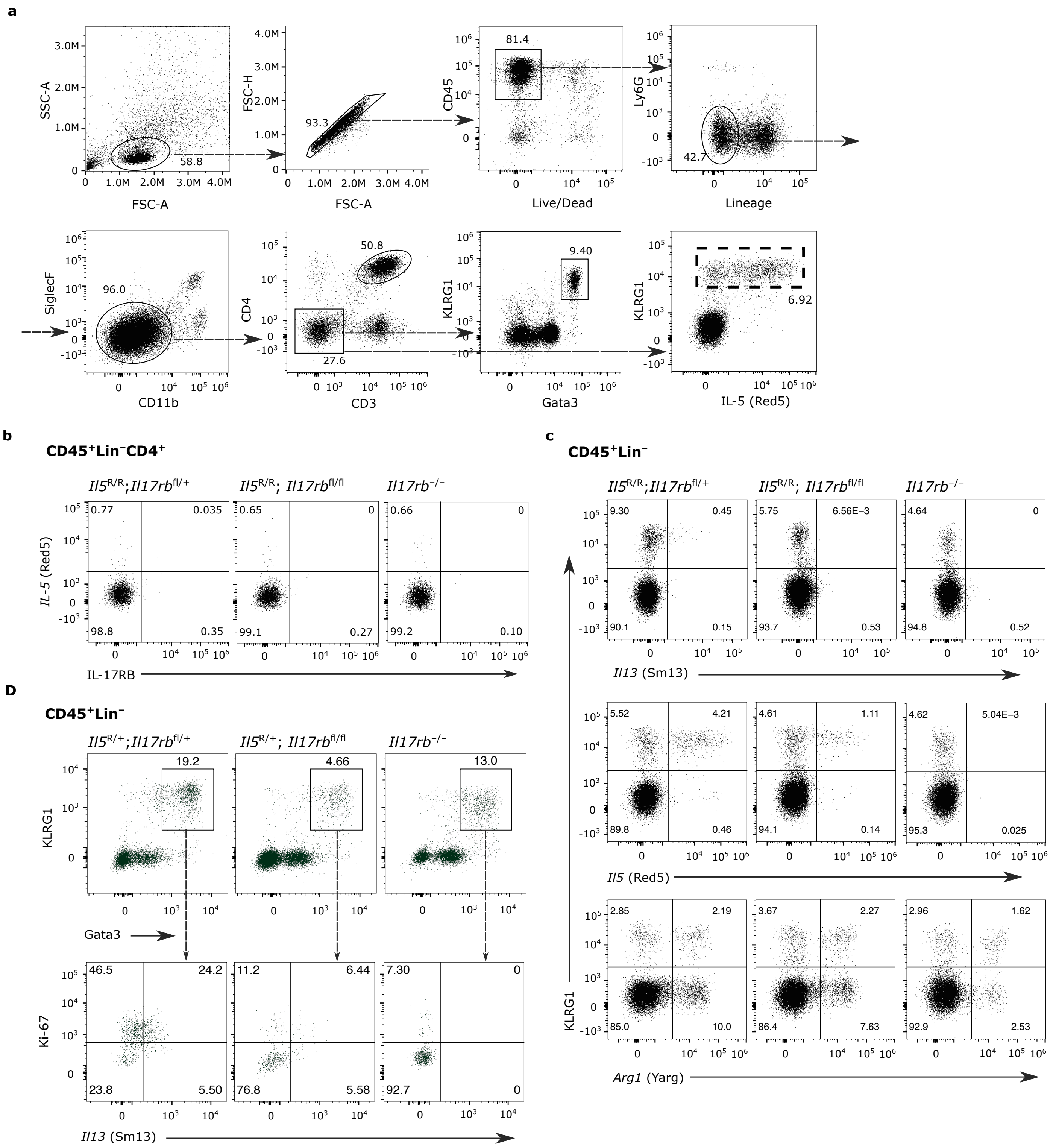
a
SSC-A
FSC-A
58.8
FSC-H
93.3
CD45
81.4
Live/Dead
Ly6G
Lineage
42.7
SiglecF
96.0
CD11b
CD4
50.8
27.6
CD3
KLRG1
9.40
Gata3
6.92
IL-5 (Red5)
b
$CD45^{+}Lin^{-}CD4^{+}$
$Il5^{R/R};Il17rb^{fl/+}$
$Il5^{R/R}; Il17rb^{fl/fl}$
$Il17rb^{-/-}$
IL-5 (Red5)
IL-17RB
c
$CD45^{+}Lin^{-}$
Il13 (Sm13)
Il5 (Red5)
Arg1 (Yarg)
KLRG1
D
$CD45^{+}Lin^{-}$
$Il5^{R/+};Il17rb^{fl/+}$
$Il5^{R/+}; Il17rb^{fl/fl}$
$Il17rb^{-/-}$
KLRG1
19.2
4.66
13.0
Gata3
Ki-67
Il13 (Sm13)

Figure 2S

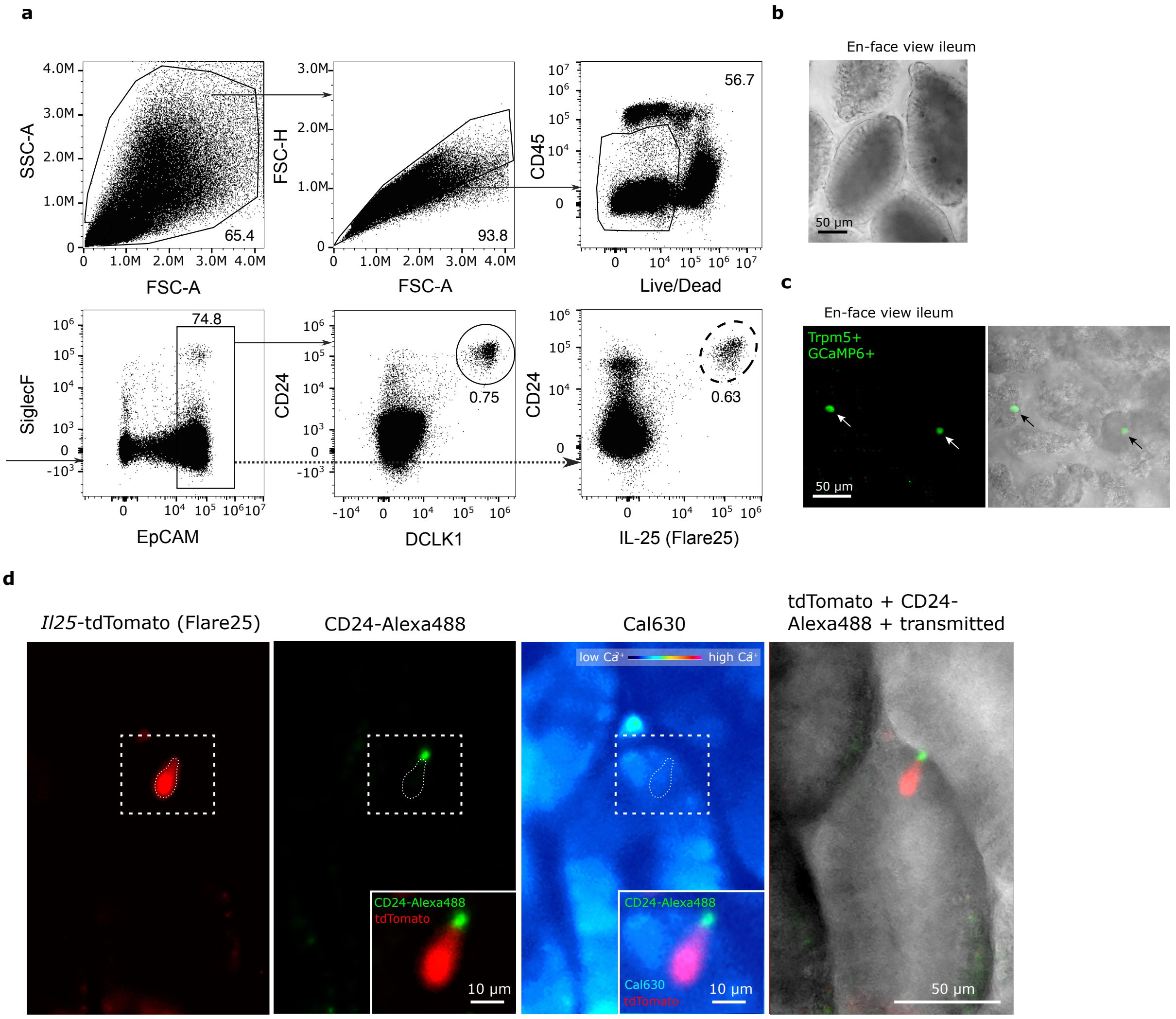

a
b
En-face view ileum
50 µm
c
En-face view ileum
Trpm5+
GCaMP6+
50 µm
SSC-A
FSC-A
65.4
FSC-H
93.8
CD45
Live/Dead
56.7
SiglecF
EpCAM
74.8
CD24
DCLK1
0.75
CD24
IL-25 (Flare25)
0.63
d
Il25-tdTomato (Flare25)
CD24-Alexa488
Cal630
tdTomato + CD24-Alexa488 + transmitted
low Ca+
high Ca+
CD24-Alexa488
tdTomato
10 µm
CD24-Alexa488
Cal630
tdTomato
10 µm
50 µm

Figure 3S

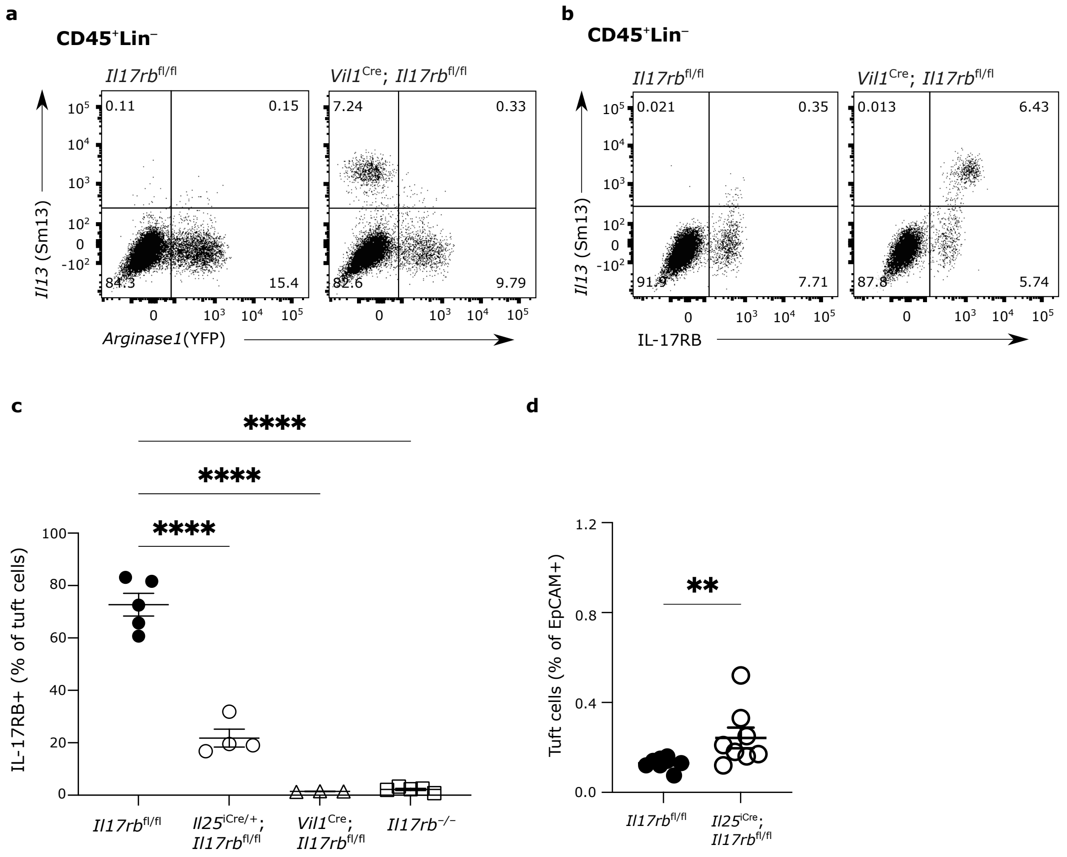

a
CD45⁺Lin⁻
Il17rb fl/fl
Vil1 Cre; Il17rb fl/fl
0.11
0.15
84.3
15.4
7.24
0.33
82.6
9.79
Il13 (Sm13)
Arginase1(YFP)
b
CD45⁺Lin⁻
Il17rb fl/fl
Vil1 Cre; Il17rb fl/fl
0.021
0.35
91.9
7.71
0.013
6.43
87.8
5.74
Il13 (Sm13)
IL-17RB
c
****
****
****
IL-17RB+ (% of tuft cells)
Il17rb fl/fl
Il25 iCre/+; Il17rb fl/fl
Vil1 Cre; Il17rb fl/fl
Il17rb −/−
d
**
Tuft cells (% of EpCAM+)
Il17rb fl/fl
Il25 iCre; Il17rb fl/fl

Figure 4S

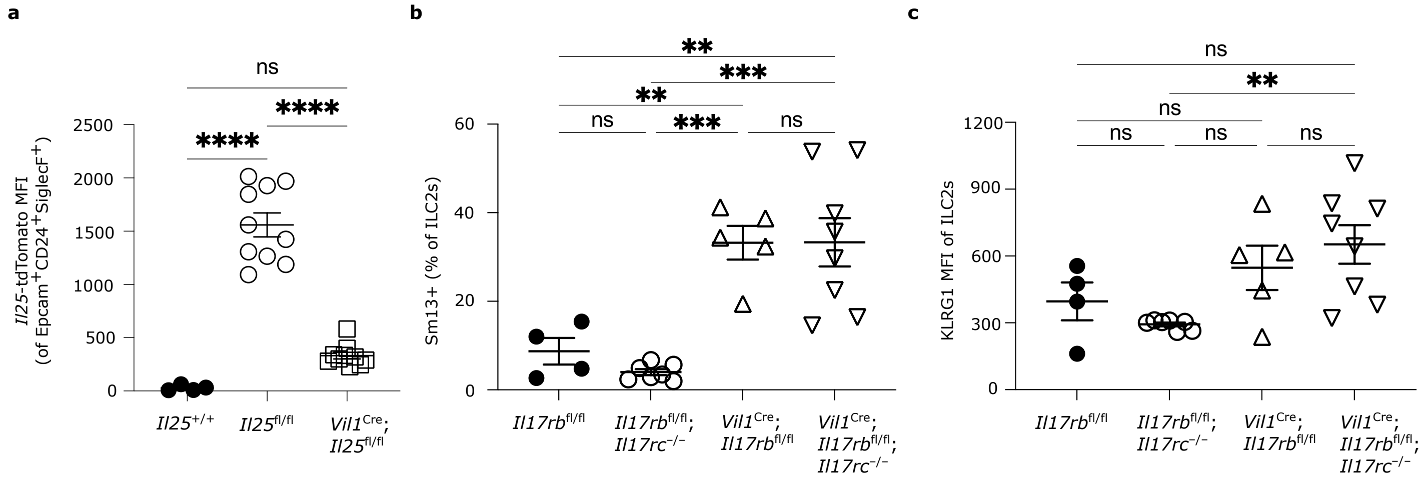
a
ns
****
****
2500
2000
1500
1000
500
0
Il25-tdTomato MFI (of Epcam+CD24+SiglecF+)
Il25+/+
Il25fl/fl
Vil1Cre; Il25fl/fl
b
**
***
**
ns
***
ns
60
40
20
0
Sm13+ (% of ILC2s)
Il17rbfl/fl
Il17rbfl/fl; Il17rc−/−
Vil1Cre; Il17rbfl/fl
Vil1Cre; Il17rbfl/fl; Il17rc−/−
c
ns
**
ns
ns
ns
ns
1200
900
600
300
0
KLRG1 MFI of ILC2s
Il17rbfl/fl
Il17rbfl/fl; Il17rc−/−
Vil1Cre; Il17rbfl/fl
Vil1Cre; Il17rbfl/fl; Il17rc−/−
